# Establishment of a Human Induced Pluripotent Stem Cell-Derived Neuromuscular Co-Culture Under Optogenetic Control

**DOI:** 10.1101/2020.04.10.036400

**Authors:** Elliot W. Swartz, Greg Shintani, Jijun Wan, Joseph S. Maffei, Sarah H. Wang, Bruce L. Miller, Leif A. Havton, Giovanni Coppola

## Abstract

The failure of the neuromuscular junction (NMJ) is a key component of degenerative neuromuscular disease, yet how NMJs degenerate in disease is unclear. Human induced pluripotent stem cells (hiPSCs) offer the ability to model disease via differentiation toward affected cell types, however, the re-creation of an *in vitro* neuromuscular system has proven challenging. Here we present a scalable, all-hiPSC-derived co-culture system composed of independently derived spinal motor neurons (MNs) and skeletal myotubes (sKM). In a model of *C9orf72*-associated disease, co-cultures form functional NMJs that can be manipulated through optical stimulation, eliciting muscle contraction and measurable calcium flux in innervated sKM. Furthermore, co-cultures grown on multi-electrode arrays (MEAs) permit the pharmacological interrogation of neuromuscular physiology. Utilization of this co-culture model as a tunable, patient-derived system may offer significant insights into NMJ formation, maturation, repair, or pathogenic mechanisms that underlie NMJ dysfunction in disease.

## Introduction

iPSCs constitute an essential tool for modeling aspects of human development and disease. Despite this, many iPSC studies on neuromuscular disorders fail to recapitulate the physiology of the NMJ by relying solely on analysis of spinal MNs (Sances et al., 2016). Evidence from patients and animal models points to dysfunction at the NMJ prior to the onset of disease symptoms, in a variety of disorders such as Amyotrophic Lateral Sclerosis (Fischer et al., 2004), Spinal Muscular Atrophy (Kong et al., 2009), and Charcot-Marie Tooth (Spaulding et al., 2016). It is therefore imperative to establish and characterize patient-specific, physiologically relevant *in vitro* models containing the essential components of the NMJ, in order to advance our understanding of these disorders (Murray et al., 2010).

Advances in hiPSC differentiation have shown formation of functional NMJs *in vitro* between hiPSC-MN and rodent sKM (Umbach et al., 2012; Yoshida et al., 2015), and between hiPSC-MNs and primary human sKM (Santhanam et al., 2018; Steinbeck et al., 2015; Afshar Bakooshli et al., 2019) or hiPSC-sKM (Puttonen et al., 2015, Osaki et al., 2018, Osaki et al., 2020). Recent demonstrations have shown that 3D neuromesodermal organoids can generate functional NMJs that consist of sKM, MNs, and Schwann cells (Faustino Martins et al., 2020).

Despite these advances, these models are ultimately difficult to scale for higher throughput assay development, as they rely on limited choice of primary human sKM, containment within a microfluidic device, or on limitations due to access of cells in a 3D structure. Moreover, useful experimental degrees of freedom are gained if all cellular components of the NMJ can be derived independently and from genetically identical and/or manipulated patient backgrounds.

Optogenetics enables the precise control of neuronal populations via light-gated ion channels (Boyden et al., 2005). Optogenetic control of the NMJ has been achieved via transgenic rodent or hiPSC-MNs paired with rodent, human primary sKM, and hiPSC-sKM *in vitro* (Steinbeck et al., 2015; Uzel et al., 2016, Osaki et al., 2018, (Afshar Bakooshli et al., 2019) and *in vivo* (Bryson et al., 2014). Optogenetics may also be paired with MEAs, enabling a system that can monitor electrophysiology of living cells in tandem with light stimulation, providing innovative applications for disease modeling or drug screening (Clements et al., 2016). Here, we leverage advances in hiPSC-differentiation, optogenetics, and MEAs to create a scalable, tunable, human patient-specific neuromuscular co-culture model system.

## Results

### Creation of stable optogenetic motor neurons

hiPSC lines were reprogrammed from fibroblasts collected from a healthy control (Line C), a *C9orf72* G_4_C_2_ expansion carrier diagnosed with behavioral variant FTD (bvFTD, Line A), and genetically corrected isogenic clones (isoA80, isoA51). All hiPSC lines were previously characterized (Swartz et al., 2016; Pribadi et al., 2016). To generate MNs, we followed a previously published protocol (Du et al., 2015) to produce populations of Hb9^+^ MNs via an adherent- or suspension-based culture protocol within 25 days, and abundant ChAT expression by day 30. (**Figure 1A-E**). We additionally generated isogenic, optogenetic reporter lines via co-transduction with two lentiviruses to enable optogenetic control of Hb9^+^ MNs (**Figure S1A**). iPSC clones were screened (see Methods, **Figure S1B-E**), resulting in selection of a single pair of stable isogenic clones (Opto-A1, Opto-isoA80-8). Suspension-based methodology yielded highly pure, Hb9^+^ MN spheres (**Figure 1F-G**), and was adopted in subsequent experiments.

**Figure 1.**
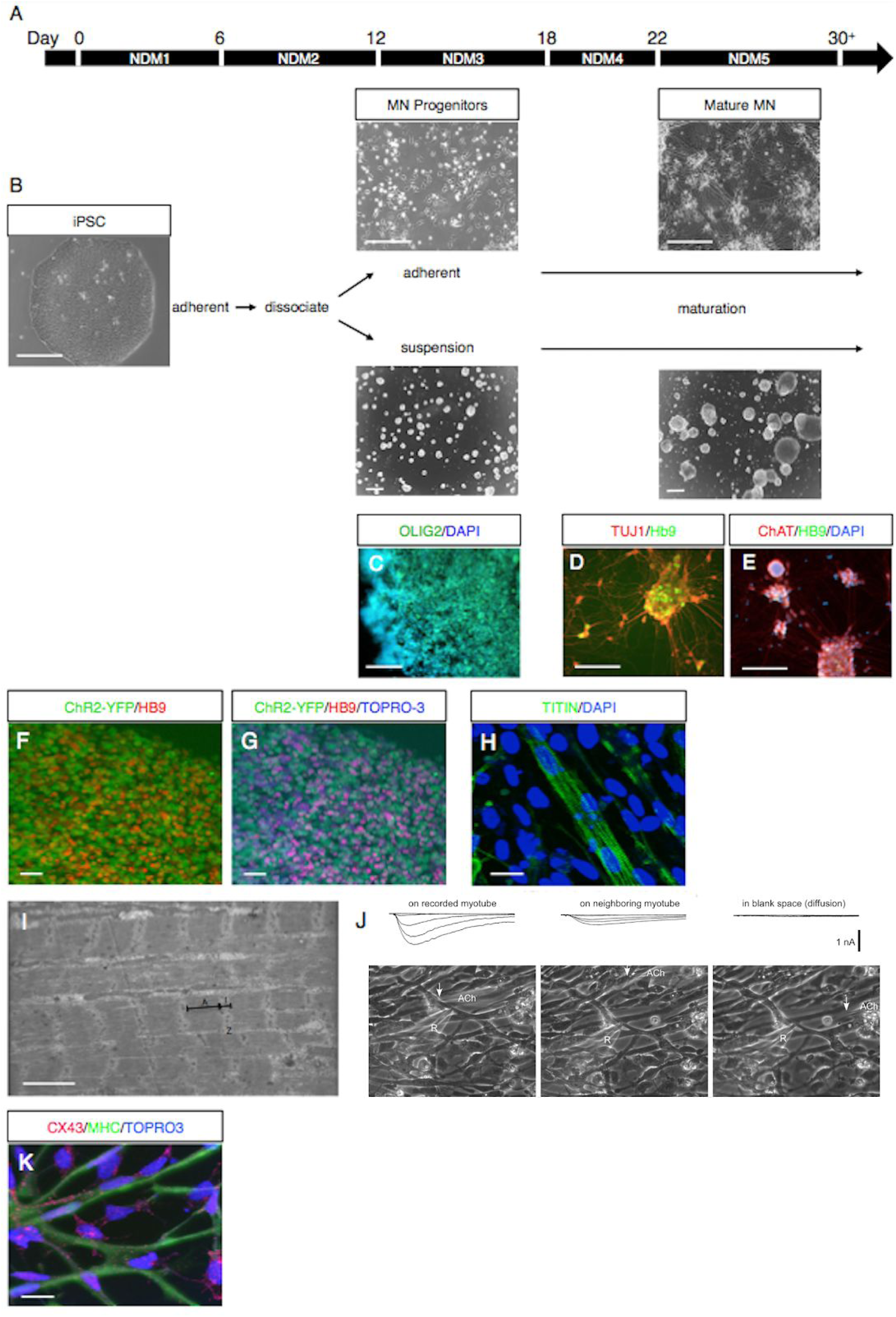
Characterization of iPSC-MN and sKM. **A**. 5-step protocol for iPSC-MNs. **B**. iPSC-MN progenitors are dissociated by day 12 and re-plated as adherent cells or grown in suspension as MN spheres. Scale bars = 250 µm. **C**. iPSC-MNs express the progenitor marker OLIG2 by day 12. Scale bar = 100 µm. **D-E**. iPSC-MNs express Hb9 by day 25 and ChAT by day 30. Scale bar = 250 µm. **F-G**. MN sphere (Opto isoA80-8) showing ChR2-YFP expression in Hb9^+^ cells. Scale bar = 20 µm. **H**. iPSC-sKM showing sarcomeric organization via Titin staining. Scale bar = 50 µm. **I**. Sarcomeric Z-lines, I-bands, and A-bands are observed in iPSC-sKM under electron microscopy. Scale bar = 1 µm. **J**. Patch clamp on iPSC-sKM shows inward current in response to puffs of acetylcholine on the myotube itself (left panel) or on the neighboring myotube (middle panel) but not when applied in blank space (right panel). **K**. iPSC-sKM cultures express Cx43 (scale bar = 10 µm).

### Spontaneous myotube contractions in 2D culture are mediated via gap junctions

We generated hiPSC-sKM following our previously published protocol (Swartz et al., 2016). Differentiated sKM showed mature sarcomeric organization by Titin staining and electron microscopy (**Figure 1H-I**), and spontaneous contractions in dense cultures. Prior to establishing co-cultures, we posited that sKM would need to be dissociated in order to distinguish NMJ-mediated contractions from spontaneous contractions. Upon dissociation and re-plating of contractile sKM, we observed a cessation in contractile activity of re-plated myotubes, although some sKM still displayed spontaneous calcium oscillations (**Figure S2A-B**). We reasoned this cessation may be due to the loss of gap junction connections, whose proteins are expressed in sKM during development (Merrifield and Laird, 2016), and were putatively observed via electron microscopy (**Figure S2C-D**).

Evidence for gap junctions in sKM was revealed by iontophoretic currents in neighboring myotubes following application of acetylcholine (**Figure 1J**), and cessation of spontaneous contractions using the reversible gap junction inhibitor 1-heptanol (**Video S1**). In addition, connexin genes expressed during myogenesis, *GJC1* (Cx45), *GJA1* (Cx43), and *GJA5* (Cx40), were also expressed in iPSC-sKM, but not in RNA from adult human sKM (**Figure S2E**). Cx43^+^ immunostaining was abundantly present in iPSC-sKM cultures (**Figure 1K**) at higher levels in the MHC^-^ cells, compared to more mature MHC^+^ myotubes, consistent with the notion that these are developmentally-regulated genes (Merrifield and Laird, 2016). Thus, spontaneous contractions observed in hiPSC-sKM are primarily mediated via gap junctions that become uncoupled during dissociation.

### Features of iPSC-derived co-cultures

Co-cultures were generated by plating MN spheres on top of dissociated sKM, as in previously reported protocols (Steinbeck et al., 2015, **Figure 2A-B**). Using control Line C, MN spheres rapidly extended axonal projections toward nearby sKM (**Figure 2C**). After 3-7 days, we verified putative NMJs via co-localization of ChAT, synaptic vesicle glycoprotein (SV2A), and the NMJ antagonist α-Bungarotoxin (α-BTX, **Figure 2D**). sKM frequently exhibited multiple sites of innervation and AchR clustering, consistent with features of NMJ development and other reports (Darabid et al., 2015; Afshar Bakooshli et al., 2019). We observed sKM hypertrophy during co-culture conditions and performed an experiment whereby co-cultures containing a single MN sphere (Opto-isoA80-8) plated on top sKM (isoA80-8) were created and the width of adjacent sKM was scored over time. sKM width significantly increased over time in the presence of a MN sphere compared with sKM grown alone in a MN- or sKM-specific medium (**Figure 2E**). This suggests that either local paracrine signals and/or electrophysiological activity derived from co-cultured MNs influence sKM hypertrophy *in vitro*.

**Figure 2.**
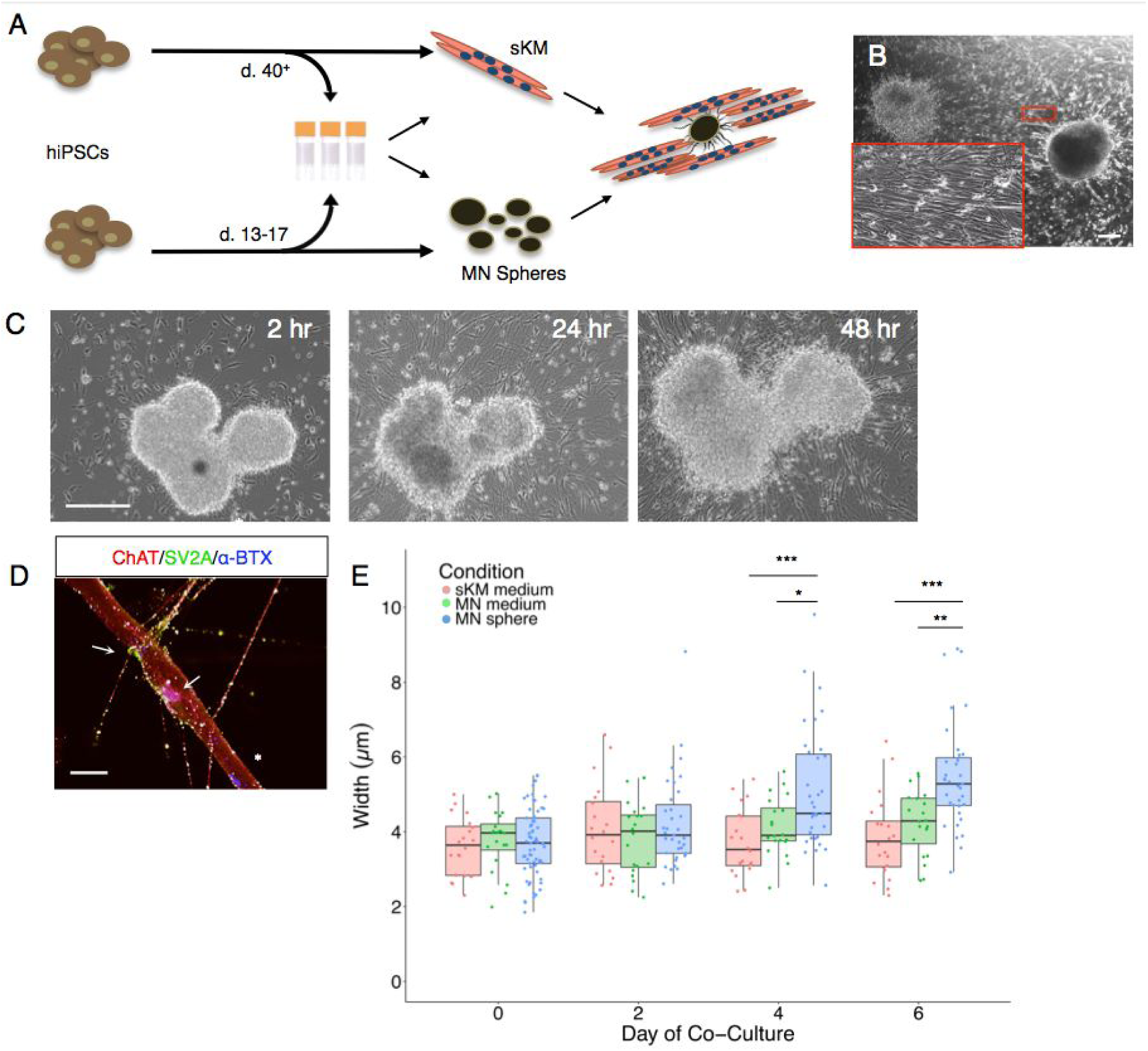
Features of iPSC-derived co-cultures. **A**. Schematic overview showing differentiation and co-culture workflow. Cells can be frozen at indicated time points and thawed to reduce time between experiments. **B**. Representative co-culture image. Inset shows sKM surrounded by axons extending radially from the nearby MN sphere. Scale bar = 250 µm. **C**. MN spheres rapidly extend projections as early as two hours post-plating and continue for at least 48 hours. Scale bar = 250 µm. **D**. A representative myotube displays multiple putative sites of innervation (arrows). Clusters of AchRs for which innervation is absent can also be seen (asterisk). ChAT antibody consistently displayed cross-reactivity with sKM. Scale bar = 20 µm. **E**. sKM undergo hypertrophy over time in the presence of a single, co-cultured MN sphere compared to sKM or MN media alone. *, p < .05; ** p < .01; *** p < .001.

### Functional co-cultures under optogenetic control

To observe if innervated sKM contained physiologically-active NMJs, we recorded spontaneous currents indicating mini end-plate potentials (mEPPs) that were subsequently abolished in the presence of the NMJ antagonist curare (**Figure 3A-C**). The majority of observed currents were under 200 pA, consistent with previous data demonstrating immediate and low-quantal acetylcholine-containing vesicle release upon muscle innervation (Chow and Poo, 1985; Sanes and Lichtman, 1999). We frequently observed contractions in adjacent sKM as early as 36 hours post co-culture. Contractions typically occurred in single myotubes but also in coordination across multiple myotubes, analogous to each myotube comprising a component of a single motor unit (**Video S2**). Thus, functional NMJs can be readily obtained in hiPSC-derived co-cultures.

**Figure 3.**
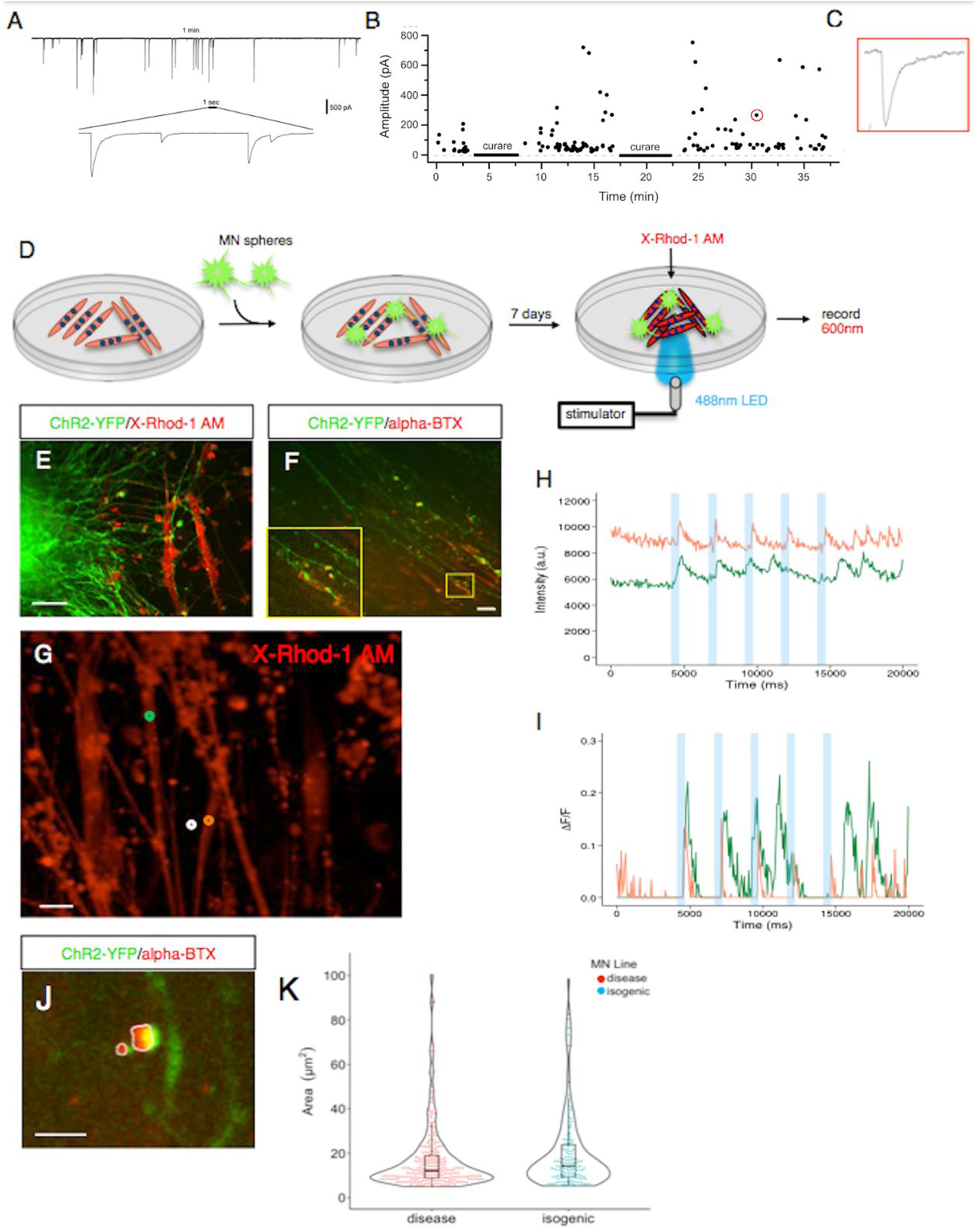
Functional co-cultures under optogenetic control. **A**. Innervated sKM display frequent, low amplitude, spontaneous bursts of current resembling mEPPs, which can be abolished by curare **(B)**. Red circle indicates a representative trace in **C. D**. Evoked calcium recording workflow, as pictured in **E**. Scale bar = 50 µm. **F**. NMJ formation by day 7 of co-culture. Scale bar = 50 µm. **G**. Innervated sKM (green and orange circles) following evoked stimulation. White circle represents the area of background normalization. Scale bar = 20 µm. **H-I**. Raw intensity (top) and background-normalized (bottom) traces of calcium flux over 20 seconds, with five 250 ms pulses of blue light. Both myotubes display spontaneous calcium flux following stimulation protocol. **J**. Method to quantify NMJ area. Scale bar = 10 µm. **K**. Isogenic MN spheres display larger NMJ areas (µm^2^) versus those from G_4_C_2_ carriers (p = .0665; effect size 2.2 (0, 4.5)).

To validate the optogenetic system, we established co-cultures of iPSC-MN spheres (Opto A1, Opto isoA80-8) and iPSC-sKM (isoA80) and recorded optogenetically-evoked calcium transients in innervated sKM using the red-shifted calcium indicator X-Rhod-1 AM (**Figure 3D-E**). We determined that 7 days of co-culture were sufficient to yield abundant NMJ formation (**Figure 3F**) coincident with reliable evoked responses in sKM. We measured evoked calcium transients in quiescent sKM, demonstrating precise sKM control across multiple light pulses (**Figure 3G-I**). Despite this success, the response in each myotube was variable and frequently resulted in spontaneous calcium flux within the sKM following stimulation, even under various stimulation protocols, making interpretation and throughput difficult for reliable phenotypic analyses.

Previous studies have shown decreased NMJ area in iPSC-derived MNs from patients with Spinal Muscular Atrophy versus healthy controls (Yoshida et al., 2015) as well as decreased NMJ number and size in *Drosophila* overexpressing *C9orf72*-associated arginine-containing dipeptides (Perry et al., 2017). Using the same isogenic lines, we scored α-BTX and YFP co-localization as a proxy for NMJ formation after 1 week in co-culture. G_4_C_2_ carrier MNs (Opto A1) displayed less total NMJ area compared to MNs derived from its isogenic control (Opto isoA80-8) (**Figure 3J-K**), suggesting that co-culture models may be useful in detection of disease- or developmentally-related NMJ phenotypes. This decrease in total NMJ area was not statistically significant, perhaps indicative of the MNs being derived from a G_4_C_2_ carrier diagnosed with bvFTD rather than ALS, or limited time in co-culture. Further studies are needed to determine if this phenotype can be recapitulated in hiPSC co-cultures.

### iPSC-derived co-cultures do not contain mature Schwann cells

*In vivo*, NMJ formation and maintenance are aided by peripheral Schwann cells that arise from the neural crest (Sanes and Lichtman, 1999). We observed at low frequency (<5%) HNK1^+^ and AP2α^+^ neural crest cells arising from our sKM protocol (**Figure S3A-I**) and HNK1^+^ (<5%) but not AP2α^+^ cells or MPZ^+^ cells from our MN protocol (**Figure S3J-K**). No MPZ^+^ cells or myelin in transverse electron microscopy sections were observed (data not shown), suggesting that neural crest cells within the co-culture do not differentiate into bona fide Schwann cells, at least in the short timeframes studied. Therefore, while functional NMJs can be obtained in co-culture, the addition of enriched Schwann cell populations may aid in further recapitulating aspects of NMJ maturation and maintenance, as demonstrated by their natural derivation in neuromuscular organoid models (Faustino Martins et al., 2020).

### Simultaneous optogenetic and electrophysiological recordings on multi-electrode arrays permit interrogation of NMJ physiology

We aimed to apply methods amenable to medium-throughput, longitudinal or drug screening studies using MEAs paired with the Lumos (Axion Biosystems) device, enabling simultaneous optogenetic and electrophysiological measurements (Clements et al., 2016). Previous studies have shown that primary rodent MN and sKM display similar extracellular action potential (EAP) characteristics (Langhammer et al., 2012; 2011). Therefore, it may be difficult to dissect cell-specific signal origin on an MEA. We first attempted to examine sKM signal origin by optogenetic control of iPSC-sKM, however, we failed to generate light-induced calcium responses after as long as 4 weeks post-transduction (data not shown) albeit while using a similar approach successfully demonstrated by others (Afshar Bakooshli et al., 2019). As an alternative strategy, we created co-cultures on multiwell MEA plates. Following 7 days of co-culture, we observed increased spikes and bursts per electrode, as well as increased network activity following light stimulation (**Video S3, Figure 4A-D**), confirming the functionality of the optogenetic system.

**Figure 4.**
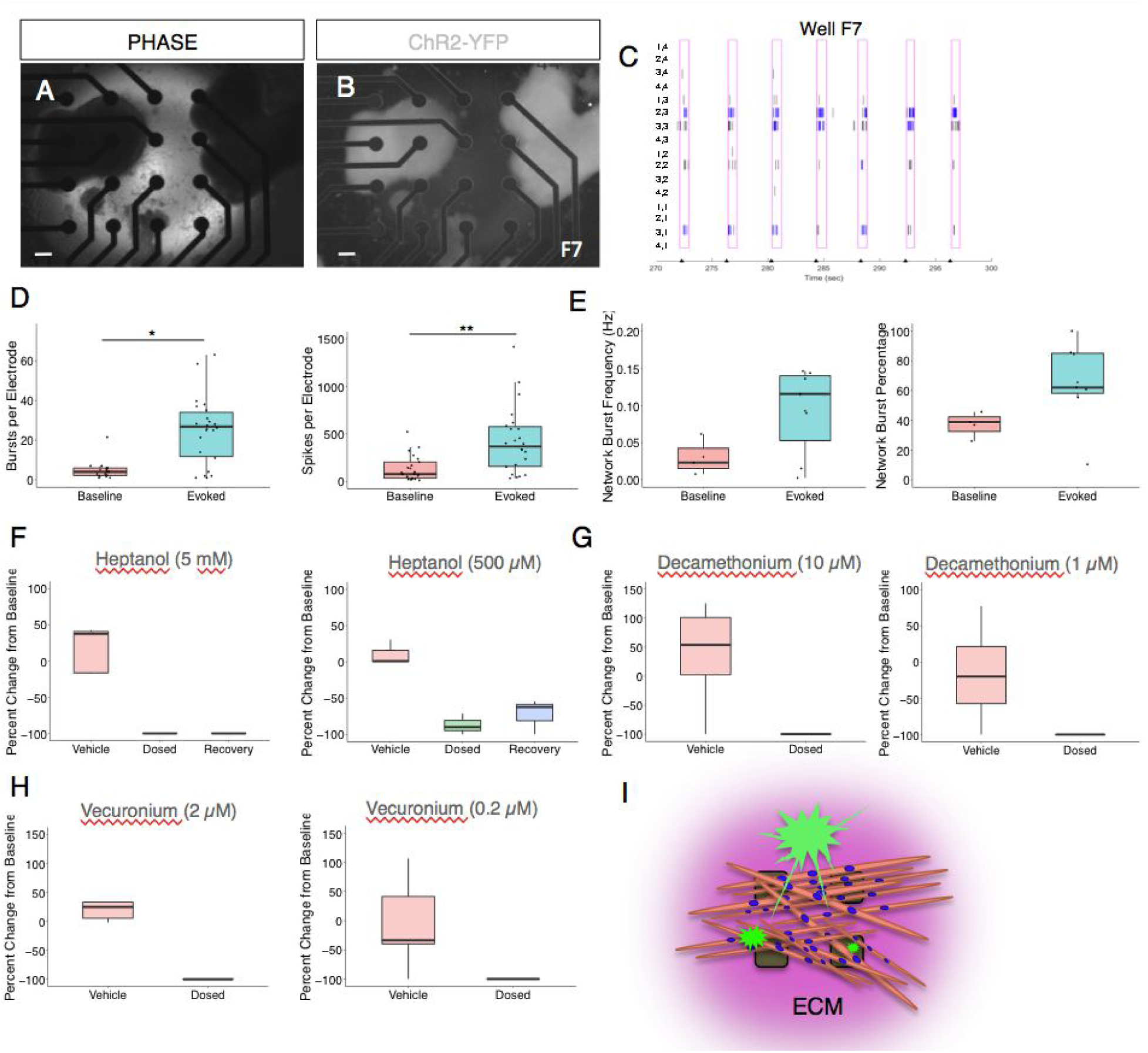
Pharmacological inhibition of evoked EAPs on MEAs. **A-B**. Co-culture on a 4×4 array of electrodes. See **Video S3**. Scale bar = 250 µm. **C**. Raster plot of the well shown in A-B during paced stimulation. Tick marks on the x-axis indicate light stimulation. Pink boxes represent evoked spike timeframe with bursts in blue. Electrode coordinates are labeled on the y-axis. **D**. Quantification of electrode activity and network activity **(E)** during baseline and stimulated recordings. Dots represent individual wells. **, p < .01; ***, p < .001. **F**. Decreased evoked spike counts following heptanol treatment, which could be partially recovered with decreased concentration or washout (recovery). **G-H**. Evoked spike counts are abolished with vecuronium and decamethonium bromide treatment. Data represented as the percent change per active electrode at baseline (untreated). **I**. Model depicting co-cultures on MEAs. Individual sKM are plated on top of protein substrate and exist in an extracellular protein matrix (ECM). MNs (green) grow on top of sKM, precluding their contact with the electrodes. In some cases, individual or small MN clusters may directly contact electrodes.

In order to parse signal origin, we treated highly responsive co-cultures with 3 drugs: the reversible gap junction inhibitor 1-heptanol, the NMJ antagonist vecuronium bromide, and the nAchR depolarization blocking agent decamethonium bromide. We quantified evoked spike counts, or those falling within a 500ms window post-stimulation, over a 5 minute recording session. Treatment with 1-heptanol led to abolishment of evoked spike counts that could be partially rescued at lower concentrations, suggesting that recorded EAPs originated from electrotonically-coupled sKM that are recorded as bursting activity, as reported by others (Rabieh et al., 2016) (**Figure 4C-E**). Support for this model was shown by heptanol treatment in C2C12 myotube cultures that yielded similar results (**Figure S4A**).

To demonstrate that recorded signals originated from functional NMJs rather than spontaneous sKM activity, we treated co-cultures with two drugs that block NMJ transmission via distinct mechanisms. Both vecuronium and decamethonium bromide completely abolished all signals, suggesting that evoked stimulation elicits NMJ transmission whereby EAPs are recorded in innervated sKM (**Figure 4E-G**). Cell morphology at active electrodes revealed dense regions of sKM lying adjacent to or underneath MN spheres (**Figure S4B-E**). Based on these data, we propose a model whereby evoked EAPs in co-culture are products of NMJ-mediated transmission originating in MNs and recorded primarily from sKM that are in contact with the electrodes (**Figure 4H**).

## Discussion

The NMJ is one of the most studied synapses in neuroscience, yet answers to fundamental questions concerning human neuromuscular development and disease are still unanswered. Our experiments demonstrate that functional, human patient-derived NMJs can be obtained *in vitro* via independent differentiation protocols to enable new avenues for investigation of basic biology and therapeutic discovery efforts adaptable to an array of neuromuscular disorders. This functionality can be controlled by light-gated ChR2 expression in innervating MNs, permitting measurement of NMJ physiology via cell permeable calcium indicators or MEAs. Thus, we have created a tunable platform for interrogating previously out of reach questions pertaining to NMJ physiology in human development or disease.

In our study, NMJs displayed features reminiscent of early NMJ development such as multiple sites of AchR clustering and innervation, lack of mature, “pretzel-like” morphology, and high-frequency but low amplitude currents indicative of low quantal mEPPs. We failed to induce robust evoked contractions in sKM during days 0-5 of co-culture, but could achieve reliable response after 7^+^ days, suggestive of NMJ maturation over time, in alignment with recent studies using human iPSC-MNs and sKM (Osaki et al., 2018, Afshar Bakooshli et al., 2019). In a similar study using optogenetic iPSC-MNs paired with primary fetal human sKM, the authors note that “contracting regions were usually identified 6 to 8 weeks” after co-culture (Steinbeck et al., 2015). Data presented in this study and others suggest that immature NMJs can be obtained in significantly shorter time frames of 1-2 weeks (Osaki et al., 2018, Afshar Bakooshli et al., 2019).

We observed difficulty in keeping co-cultures stable for over 10 days, similar to other reports (Umbach et al., 2012). During co-culture, we observed axonal retraction over time that may be analogous to activity-dependent synaptic elimination at the NMJ (Darabid et al., 2015) and further studies utilizing tagged NMJ proteins such as AchR or Rapsyn in combination with time-lapse imaging may provide answers to these phenomena. Alternatively, given that we did not observe bona fide Schwann cells in our co-culture, the addition of independently derived Schwann cells may stabilize the co-culture for longer-term studies and enable further NMJ maturation. Although sKM displayed hypertrophy during co-culture, use of bioengineering strategies such as compartmentalized microfluidic devices or nano-patterned hydrogels may also provide a more native environment for each respective cell type, leading to stable, long-term co-cultures (Ionescu et al., 2016; Langhammer et al., 2012; Santhanam et al., 2018; Uzel et al., 2016; Zahavi et al., 2015, Osaki et al., 2018).

Previous studies have shown co-culture recordings on MEAs, however this required custom-made devices to physically separate cell types (Langhammer et al., 2012). To validate functionality and determine cell-type specific signal origin, we utilized optogenetics and drugs with known pharmacology. This revealed that recorded signals likely originated primarily from electrotonically-coupled sKM, rather than MNs, which may have otherwise been expected. These findings therefore support the use of MEAs as a tool for medium-throughput screening of small molecules or growth factors for their effects on patient-specific NMJ physiology.

## Acknowledgements

This work was supported by the Adelson Medical Research Foundation and the Tau Consortium. We would like to thank Marianne Cilluffo (UCLA) for assistance in preparation of electron microscopy samples, Emily Matheu and Lucy Chammas (Axion Biosystems) for assistance with multi-electrode array data collection, and Dr. Ken Yamauchi and Dr. Bennett Novitch (UCLA) for lending lentiviral optogenetic constructs and training on microscopy equipment.

## Author Contributions

Conceptualization, E.W.S. and G.C; Methodology, E.W.S., G.S.; Validation, E.W.S., G.S., S.W.; Formal Analysis, E.W.S. and J.M.; Investigation, E.W.S., G.S., and J.W.; Writing – Original Draft, E.W.S. and G.C., Visualization, E.W.S.; Supervision, L.H. and G.C.; Funding Acquisition, G.C., B.L.M.

## Abridged Experimental Procedures

Full materials and methods are described in the Supplemental Experimental Procedures.

### iPSC Differentiation and Co-Culture Assembly

iPSCs, iPSC-sKM (Swartz et al., 2016), and iPSC-MNs (Du et al., 2015) were grown and differentiated as done previously, with minor variation. Co-cultures were assembled using dissociated iPSC-sKM between days 60-100 *in vitro*, followed by iPSC-MN spheres between days 40-60 plated on top, following Steinbeck et al., 2016, with variations. Co-cultures were grown in BrainPhys containing neurotrophic factors and recombinant human Agrin (50 ng/mL). Co-cultures were kept for experimentation up to 2 weeks.

### Creation of Stable, Optogenetic iPSC lines

Isogenic *C9orf72* lines were created as previously described (Pribadi et al., 2016). iPSCs were stably transduced with two lentiviral constructs and screened. 1 clone from each line with the highest transgene expression was chosen for further study.

### Myotube Width and Neuromuscular Junction Area Calculations

Co-cultures were created using identical batches of sKM and MNs. Co-cultures were grown in 3 conditions: (1) in sKM medium, (2) in MN medium, or (3) in MN medium with a single MN sphere. sKM were imaged every 48 hours of co-culture until day 6. Images were scored manually across >5 individual coverslips per condition. On day 7 of co-culture, condition 3 cells were fixed, stained with α-BTX (1:500), and imaged. Co-localized α-BTX and YFP plaques were scored by a blinded observer. Areas below 5 µm^2^ were excluded from analysis as potentially indistinguishable from background.

### Optogenetically-evoked Calcium Recordings

Co-cultures were incubated with the cell permeable calcium dye X-Rhod-1 AM (2µM) prior to imaging on a Zeiss Axio Observer Z1 spinning disc confocal microscope outfit with an EMCCD camera, incubation system, and blue LED (ThorLabs, M470L02, 470nm). Stimulation settings were controlled by a Master-8 stimulator. Data was collected using ZENBlue software (Zeiss) in 20 second videos where the stimulation paradigm consisted of five 250 ms LED pulses at 1 Hz. Regions of interest (ROIs) were manually selected on individual myotubes and normalized to a background ROI containing the same pixel volume (4.5 pixel diameter circle). ΔF/F ratiometric measurements were performed using ZENBlue software.

### Multi-Electrode Arrays

Co-cultures were grown for 7 days prior to recording. Electrophysiological activity was recorded using the Maestro Pro (Axion Biosystems). Data was acquired as raw voltage using a bandpass filter (200 – 4000 Hz) and a gain of 1000x at a sampling rate of 12.5 kHz. Neural spikes were identified as peaks in voltage surpassing 6x the standard deviation on each electrode. Optogenetic stimulation was performed with the Lumos (Axion Biosystems) using 475 nm light at 50% power with 500 ms pulses at 0.25 Hz. Prior to drug treatment, baseline evoked activity was recorded for 5 minutes, followed by 20 minute incubation in vehicle and recording. This protocol was then repeated for each drug treatment, using the same well that previously received vehicle.

Gross functional activity was quantified as the number of spikes per electrode during recording. Synchronous well-wide evoked activity was quantified as the network burst frequency and the percentage of spikes that belong to a network burst. Response to drug treatment in optically-paced cultures was quantified by the average number of spikes in a 500 ms window post-stimulation only on electrodes yielding an evoked response.

## Supplementary Figures

**Figure S1.**
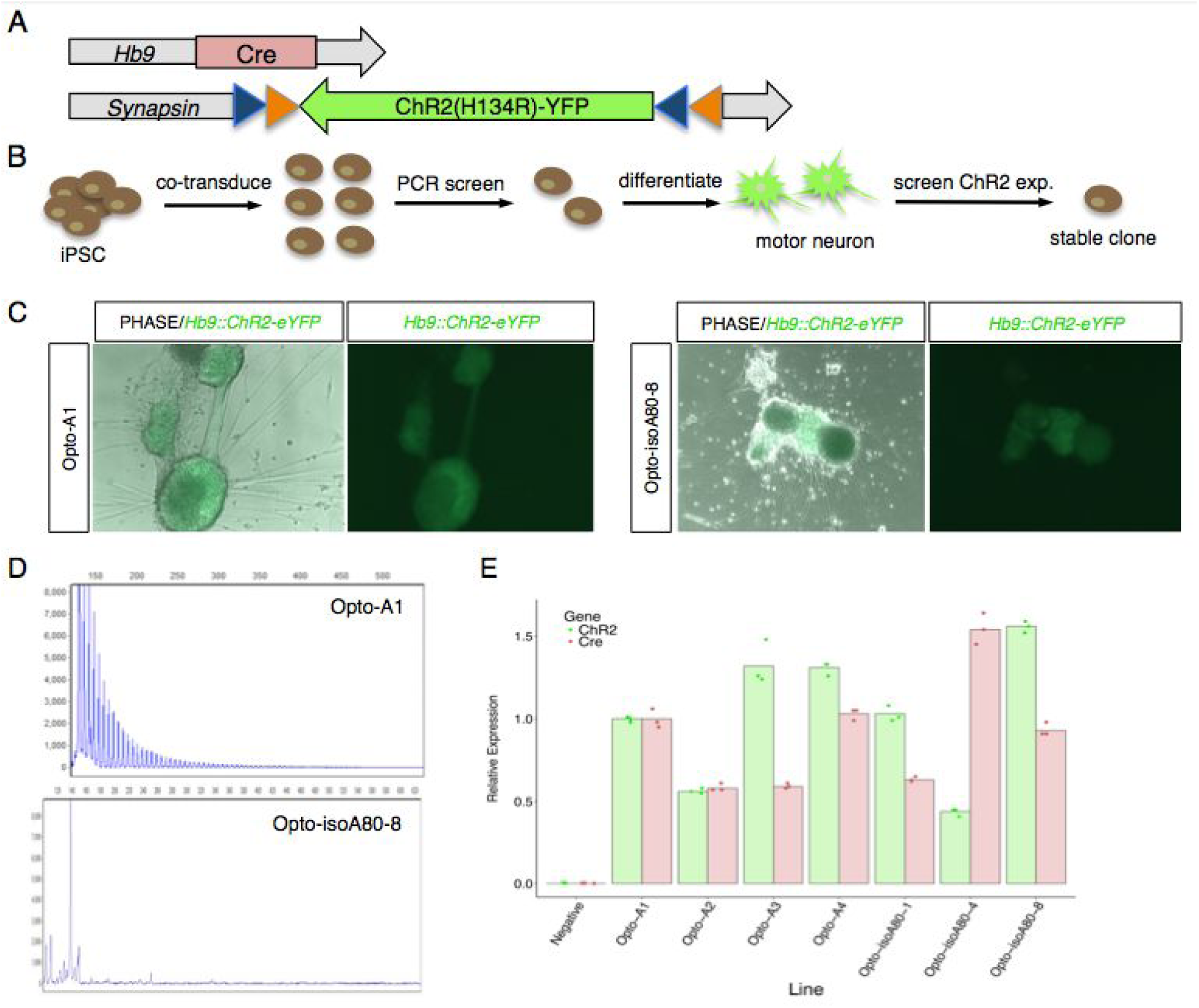
Generation of stable optogenetic iPSC lines. **A**. Schematic of two lentiviral vectors used to generate optogenetic lines. When Cre is expressed in Hb9^+^ cells, it produces serial recombination at independent *loxP* (orange triangles) and *lox2722* (blue triangles) sites, leading to the ChR2-YFP fusion construct expressed in the correct reading frame. **B**. Schematic representing the screening and selection process for isolation of transgenic iPSC clones. **C**. Representative images of ChR2 expression in isogenic iPSC-MN lines at day 30 of differentiation. Scale bar = 250 µm. **D**. Repeat-primed PCR confirmed the G_4_C_2_ status of the selected optogenetic iPSC clones. Cells positive for the expansion display a stereotypical sawtooth pattern (top). **E**. qPCR of *Cre* and *ChR2* expression in selected iPSC-MN clones at day 30 of differentiation. Opto-A1 and Opto-isoA80-8 were chosen for further study. Negative represents non-transduced iPSCs. Data are normalized to the geometric mean of housekeeping genes *GAPDH* and *ACTB*.

**Figure S2.**
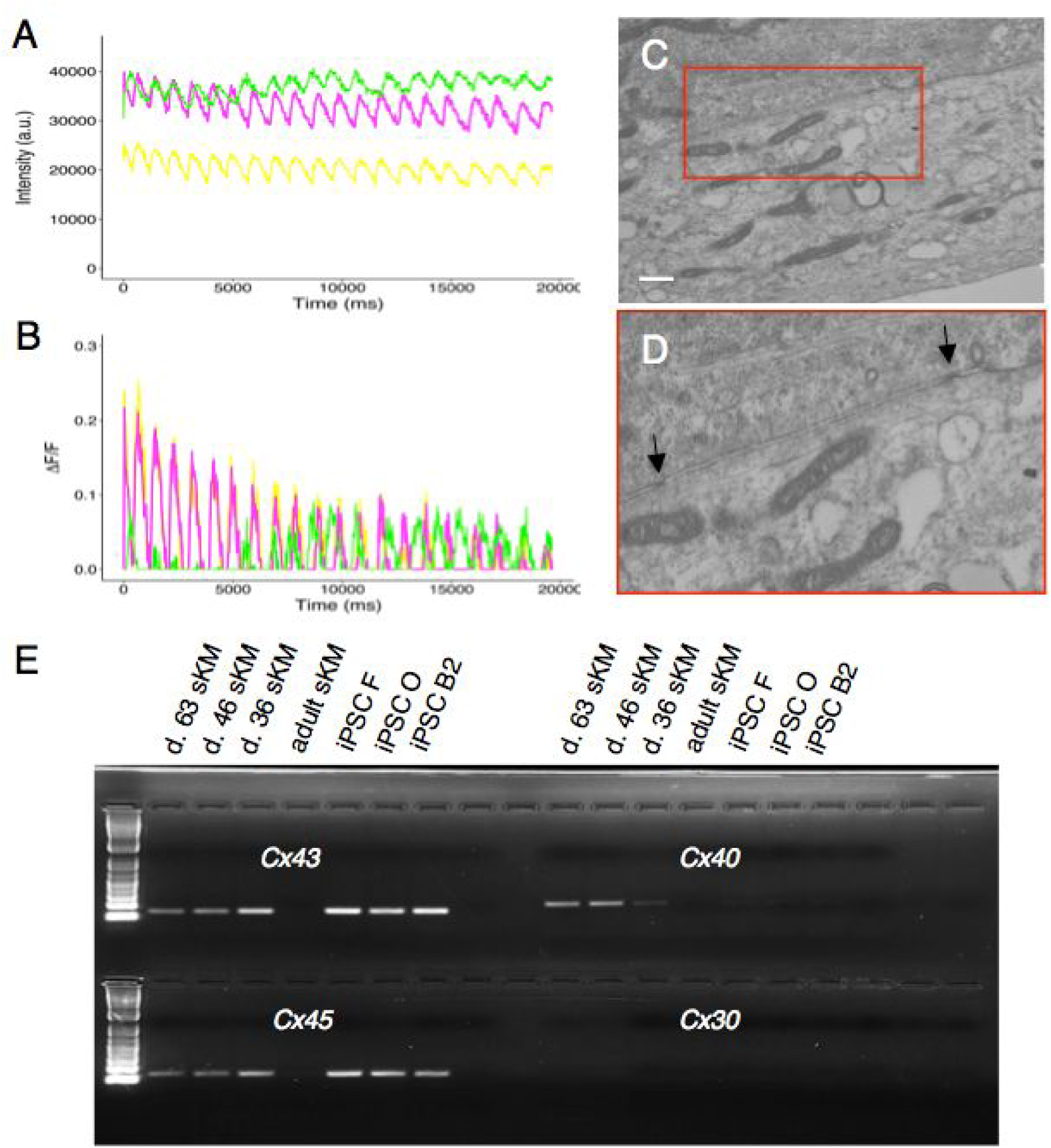
Gap junctions in iPSC-sKM. **A-B**. Representative raw and background-corrected traces of spontaneous calcium flux without contraction in dissociated sKM. **C-D**. Electron microscopy displays putative gap junction connections (arrows) in adjacent myotubes. Scale bar = 1 µm. **E**. iPSC-sKM (line isoA80) but not adult human skeletal muscle expresses all 3 predicted gap junction genes, *Cx43, Cx40, and Cx45*, at 3 timepoints of differentiation. 3 undifferentiated iPSC lines, which are known to express *Cx43* and *Cx45* but not *Cx40* were run as positive controls. The cochlear-specific gene *Cx30* served as a negative control.

**Figure S3.**
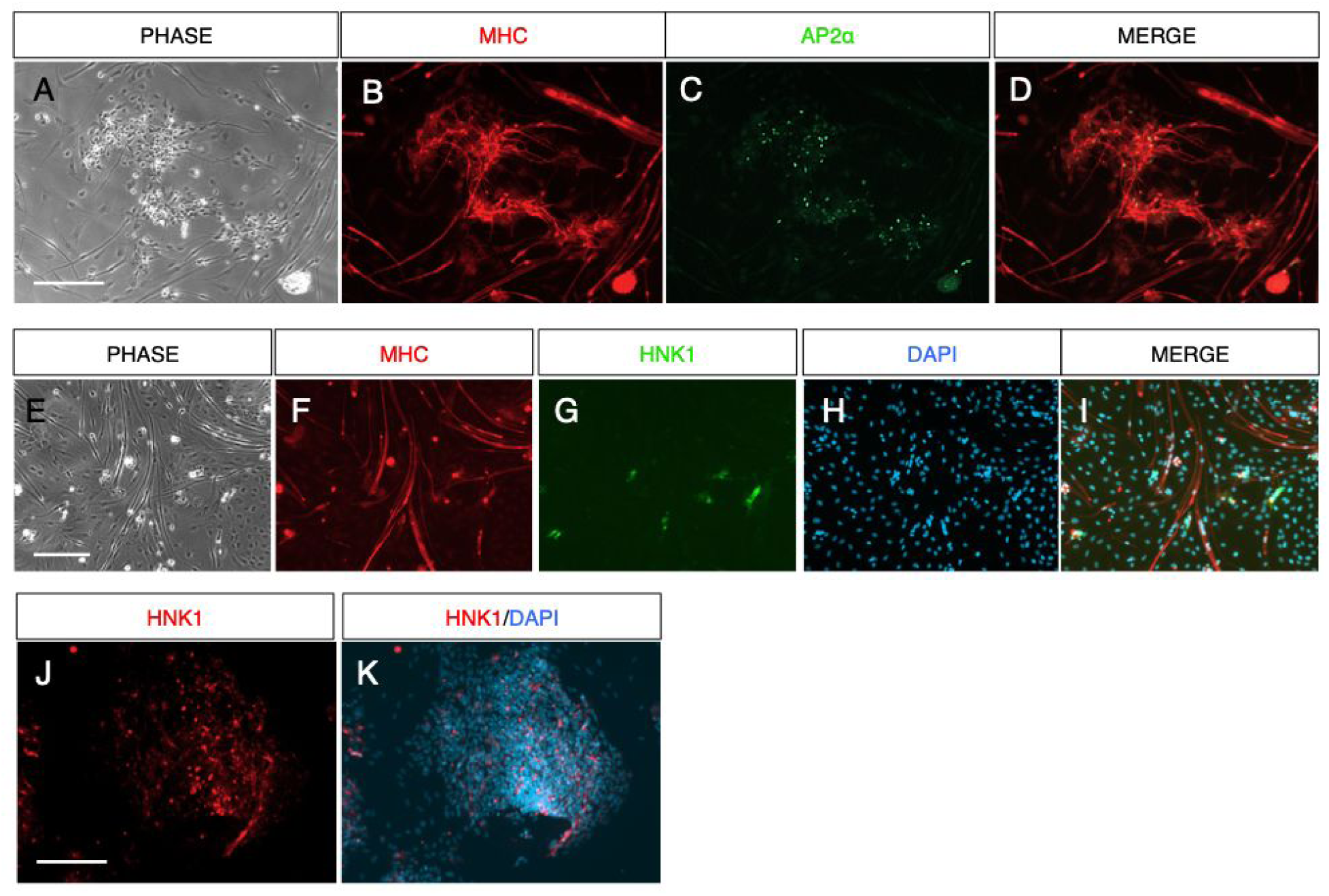
Neural crest cells are present in iPSC-sKM and MN sphere culture protocols. **A-D**. AP2α^+^ cells are present in low frequency at non-MHC^+^ differentiated regions of iPSC-sKM cultures. MHC antibody consistently displayed cross-reactivity with neuronal or neural crest cell types with small somas. **E-I**. HNK1^+^ cells are present in low frequency at non-MHC^+^ differentiated regions of iPSC-SKM cultures. **J-K**. HNK1^+^ but not AP2α^+^ cells were detected in non-specific differentiated regions of iPSC-MN cultures.

**Figure S4.**
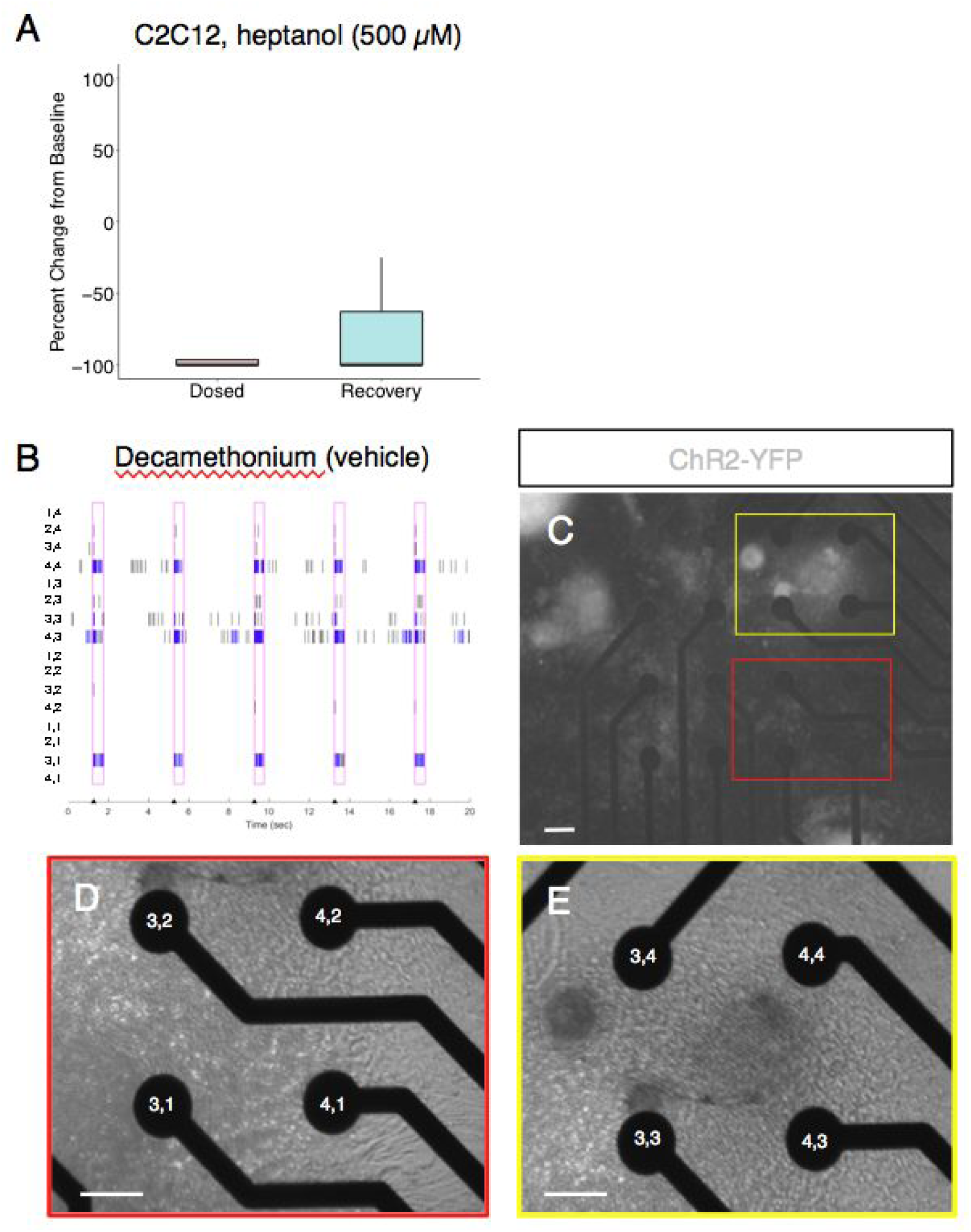
Electrode level morphology analysis. **A**. Spike counts decrease following heptanol treatment in C2C12 myotubes. **B**. Representative raster plot during recording from a vehicle-treated well which later received decamethonium bromide treatment, shown in **Figure 4G**. y-axis represents electrode coordinates. **C**. View of motor neuron location in the well recorded in **B**. Black circles represent individual electrodes. Scale bar = 250 µm. **D-E**. Morphology at active electrodes (4,4; 3,3; 4,3; and 3,1) reveals dense myotube morphology in close proximity to motor neurons. Scale bar = 250 µm.

**Video S1: iPSC-sKM spontaneous contractions are mediated by gap junctions**.

iPSC-sKM contractile activity is abolished following treatment with the reversible gap junction inhibitor 1-heptanol. Highly contractile myotubes were recorded in 20 second videos at baseline and time = 5 and 10 minutes following bath application of 5 mM 1-Heptanol. Contractile activity returned to a baseline level 15 minutes after washout. Myotubes were at day 48 *in vitro* at the time of recording.

**Video S2: iPSC-sKM contractions in co-culture**.

iPSC-MN spheres provide synaptic input to multiple dissociated iPSC-sKM, which elicit single or coordinated spontaneous contractions.

**Video S3: Optogenetic pacing of co-cultures on multi-electrode arrays**.

Co-cultures were prepared across all 48 wells and baseline spontaneous activity was recorded. Firing activity in wells became synchronized, with some instances of intermediate spontaneous firing, following blue light optogenetic pacing at 0.25 Hz. Video displayed at 2x speed. Well F7 and plate-wide data are captured in **Figure 4A-E**.

## Supplemental Experimental Procedures

### iPSC Maintenance

Fibroblasts were obtained from skin biopsies of patients carrying the *C9orf72* G_4_C_2_ expansion and healthy controls from the UCSF Memory and Aging Center with patient consent and Institutional Review Board approval (#10-000574). Fibroblasts were cultured in DMEM + 10% Fetal Bovine Serum (FBS). Reprogramming was performed with the STEMCCA lentiviral vector containing *OCT4*, c-*MYC, SOX2*, and *KLF4* (Millipore). Pluripotency was assessed via Pluritest (Müller et al., 2011) and immunocytochemical staining of NANOG, OCT4, SSEA4, and TRA-181 (Swartz et al., 2016). Karyotyping was performed by Cell Line Genetics between passages 5-25 following reprogramming. iPSCs were cultured at 37°C and 5% CO_2_ on Vitronectin (Stem Cell Technologies) coated plates in TeSR-E8 media (Stem Cell Technologies) and passaged every 4-6 days using the ReLeSR reagent (Stem Cell Technologies) following the manufacturer’s instructions. iPSC lines used in this study and relevant patient information is listed below in **Table S1**.

**Table S1:** Patient information for iPSC lines used in the study. bvFTD: behavioral variant FTD; Y.O.B: year of birth; exp, G_4_C_2_ repeat expansion. “Opto” indicates genetically integrated viral vectors that enable optogenetic experiments.

| Parent Cell Line | Line Name | Status | C9orf72 Genotype | Diagnosis | Gender | Race | Y.O.B |
| --- | --- | --- | --- | --- | --- | --- | --- |
| C1B1 | C | Control | (+/+) | Unaffected | Male | White | 1945 |
| CC3A4 | B2 | Carrier | (+/exp) | FTD/ALS | Male | White | 1963 |
| F7A4 | F | Control | (+/+) | Unaffected | Male | Unknown | 1968 |
| O1A1 | O | Sporadic | unknown | bvFTD | Male | White | 1941 |
| OO6A2 | A | Carrier | (+/exp) | bvFTD | Male | White | 1965 |
| OO6A2 | Opto-A1 | Carrier | (+/exp) | bvFTD | Male | White | 1965 |
| OO6A2 | isoA51 | Isogenic Control | (+/Δ) | bvFTD | Male | White | 1965 |
| OO6A2 | isoA80 | Isogenic Control | (+/$\Delta$) | bvFTD | Male | White | 1965 |
| OO6A2 | Opto-iso A80-8 | Isogenic Control | (+/$\Delta$) | bvFTD | Male | White | 1965 |

## Morphogen-Directed iPSC Differentiation

### Motor Neuron Differentiation

For differentiation into spinal motor neurons, iPSCs were differentiated according to previous reports, with some variations (Du et al., 2015). For co-culture experiments, cells were grown using a hybrid process of adherent and suspension culture, resulting in MN spheres. For immunocytochemistry, cells were grown adherent throughout, with a passaging step between days 12-18 of differentiation onto 96 well plates coated in Poly-L-Ornithine overnight followed by 3 washes in sterile water followed by 1 hour of drying. The plates were then coated with Laminin (15 µg/mL, ThermoFisher) and incubated at 37°C for 1 hour prior to the addition of cells. The following basal medium formulation was used throughout the differentiation process: A 1:1 mix of basal mediums DMEM/F12 and Neurobasal supplemented with 1% Sodium Pyruvate, 1% Non-Essential Amino Acids, 1% GlutaMAX, 0.5% N2 supplement, and 0.5% B27 supplement. Approximately 24-48 hours after passaging, small iPSC colonies were switched into a neural differentiation medium (NDM1) consisting of the base medium supplemented with ascorbic acid (100 ng/mL), SB43152 (10 µM), LDN193189 (200 nM), and CHIR99021 (3 µM) for 6 days, changing medium every other day or every day as needed. On day 6, the medium was switched to NDM2 consisting of the base medium supplemented with ascorbic acid (100 ng/mL), cAMP (0.5 µM), SB431542 (10 µM), LDN193189 (200 nM), CHIR99021 (1 µM), retinoic acid (200 nM), and purmorphamine (0.5 µM) for an additional 6 days, changing ¾ medium every day. On day 12, the medium was switched to NDM3 consisting of the base medium supplemented with ascorbic acid (200 ng/mL), cAMP (0.5 µM), retinoic acid (500 nM), and purmorphamine (0.1 µM) for an additional 6 days, changing ¾ medium every day.

For co-culture experiments, cells were dissociated with Accutase (ThermoFisher) between days 13-17, re-plated in ultra-low attachment plates (Corning), and grown as MN spheres in suspension. MN spheres were passaged as necessary by gentle dissociation following 8-12 minutes of Accutase treatment, followed by gentle pipetting and re-plating in low attachment plates. For immunocytochemistry experiments, cells were gently dissociated with Accutase between days 14-17 and re-plated on 96 well plates previously coated with poly-L-ornithine (Sigma) for 24 hours overnight followed by 3 washes in sterile water followed by 1 hour of drying. The next day, the plates were coated with Laminin for 1 hour at 37°C prior to adding cells. In each case, motor neuron progenitors (MNPs) were frozen down between days 14-17 (see Cryopreservation of iPSC-derived Cells).

On day 18, cells were switched to NDM4 consisting of the basal medium supplemented with ascorbic acid (200 ng/mL), cAMP (1 µM), DAPT (10 µM), retinoic acid (500 nM), and purmorphamine (0.1 µM) for an additional 4 days, adding 0.75-1.0 mL of medium every other day for suspension culture or replacing ¾ medium every other day for adherent culture. On days 22 onward, cells were cultured in NDM5 consisting of the base medium supplemented with ascorbic acid (200 ng/mL), cAMP (1 µM), BDNF (10 ng/mL), GDNF (10 ng/mL), and CNTF (10 ng/mL), adding 0.75-1.0 mL of medium every other day. To avoid aggregation, MN spheres were collected once per week in 15 mL conical tubes and allowed to pellet by gravity for 5-10 minutes. Single cells in solution were aspirated and MN spheres were then broken apart using gentle pipetting with a P200 pipette tip or mechanical agitation and re-plated in low attachment plates. Motor neuron spheres were cultured until ∼day 40-50 for use in downstream experiments and adherent motor neurons were cultured until day 30 until fixation for immunocytochemical staining.

### Skeletal Muscle Differentiation

iPSC-sKM were generated as previously described with minor variation (Swartz et al., 2016). 1.5% DMSO in TeSR-E8 medium was added when iPSC colonies were roughly 250-400 µm in diameter. On day 0, cells were cultured in CDM FLyBC for **48** hours (changed from 36 hours). Chemically defined media (CDM) consisted of a 1:1 mixture of IMDM and Ham’s F12 media with the addition of Bovine Serum Albumin (5 mg/mL), 100X Lipids (1X), Transferrin (15 µg/mL), 1-Thioglycerol (450 µM), and Insulin (7 µg/mL). FLyBC consists of FGF2 (20 ng/mL), LY294002 (10 µM), BMP4 (10 ng/mL), and CHIR99021 (10 µM). After **48** hours (changed from 36 hours), cells were cultured in CDM FLy for an additional 5.5 days. On day 7, cells were cultured in MB-1 and 15% FBS (3/4 daily media change), generously donated from the laboratory of Dr. Jacques Tremblay (Université Laval, Canada) and prepared per their instructions. On day 12, cells were cultured in Fusion medium (replaced alternating days) consisting of 2% Horse Serum in DMEM. After 23 days *in vitro*, mature myotubes were cultured in N2 medium, composed of DMEM/F12 supplemented with 1% Insulin-Transferrin-Selenium solution and 1% N2 supplement, until a majority of the cells displayed spontaneous contractions or sufficient hypertrophy based on qualitative assessment, typically between days 40-70 *in vitro*. Myotubes were dissociated completely in Accutase for ∼25 minutes and either used directly in co-culture experiments or frozen in cryopreservation medium. Myotubes were then thawed as needed for co-culture experiments.

For C2C12 cells, low passage myoblasts were grown on 0.1% gelatin in DMEM and 10% FBS until cells were approximately 90% confluent. At this stage, myoblasts were switched to a fusion medium consisting of DMEM and 2% horse serum and cultured for 7^+^ days or until myotubes were abundant. Myotubes were kept in fusion medium, changing ¾ medium every 3 days, until experimentation.

### Assembly of Co-Culture

Co-cultures were performed on sterilized glass coverslips within 12 well plates or glass bottom microwell dishes (MatTek). Coverslips or dishes were coated in Matrigel, incubated for 1-2 hours at 37°C, and aspirated. For coverslip preparation, a P20 pipette was used to move the coverslip within the middle of the well, avoiding contact with the edge of the well. All myotube and motor neuron plating was performed qualitatively without cell counting. A concentrated solution of skeletal myotubes taken directly from a dissociated live culture or freshly thawed vial was prepared in N2 medium. 20-40 µL of cells were added drop-wise to the coverslip or glass bottom dish, covering the surface evenly. The amount of cells added was qualitatively assessed under a microscope until a sufficient amount of cells were plated, aiming for 40-50% confluence. Additional N2 medium was added in a sufficient volume to maintain surface tension on the glass surface. The cells were then incubated overnight.

The next day, myotubes were rinsed once in N2 medium and then cultured in N2 medium for an additional 3-4 days to allow for hypertrophy and/or proliferation. After this time period, a 1:1 mixture of NDM5 and NDM6 were prepared to avoid osmotic shock. NDM6 consisted of a basal medium of BrainPhys supplemented with 1% Sodium Pyruvate, 1% Non-Essential Amino Acids, 1% GlutaMAX, 1% N2-A supplement, 2% SM1 supplement, ascorbic acid (200 ng/mL), cAMP (1 µM), BDNF (10 ng/mL), GDNF (10 ng/mL), and CNTF (10 ng/mL). MN spheres were pelleted by gravity for 5-10 minutes and re-suspended in a low volume of 1:1 NDM5 and NDM6 containing a 1:50 dilution of Geltrex (ThermoFisher). N2 medium was aspirated from the coverslips or glass bottom dish. Using a P20 pipette, 20 µL of MN spheres were added drop-wise to the myotubes, aiming for multiple visible MN spheres plated per coverslip or dish. An additional volume of 1:1 NDM5 and NDM6 containing a 1:50 dilution of Geltrex was added in a sufficient volume to maintain surface tension. The cells were then incubated overnight.

The next day, cells were gently rinsed with a 1:1 mixture of NDM5 and NDM6 and then cultured in a 1:1 mixture of NDM5 and NDM6 containing recombinant human neuronal Z-isoform Agrin (50 ng/mL, R&D Systems). A ¾ medium change was performed every 2-3 days with NDM6 containing recombinant human Agrin (50 ng/mL). In some cases, the mitotic inhibitor cytosine arabinoside (Ara-C, 5 µM, Sigma) was added for 48 hours to prevent proliferation of undefined progenitor cell types. Co-cultures were kept for experimentation up to 2 weeks.

A complete list of all medium formulations and timeframes of use can be found below in **Table S2**.

**Table S2.**
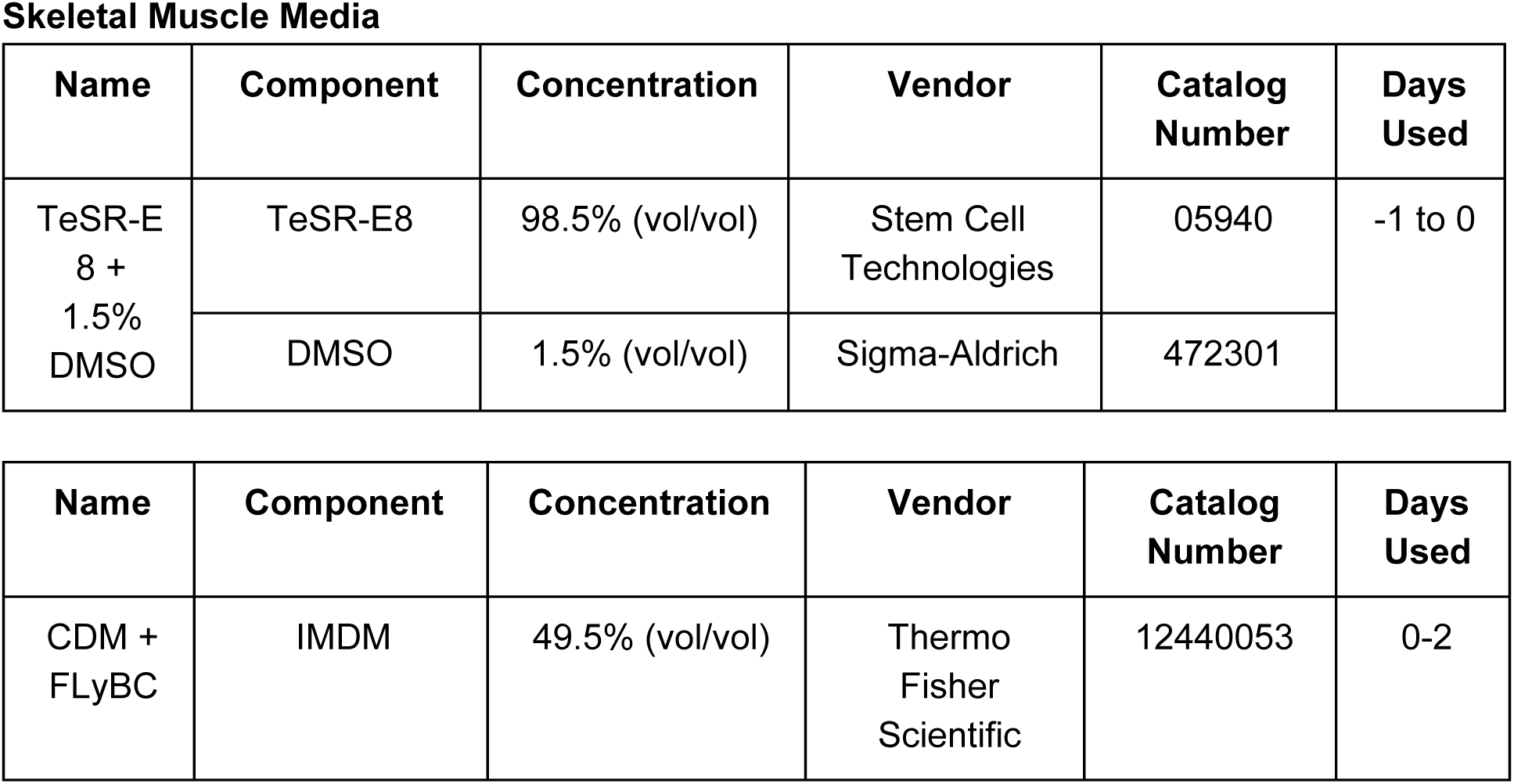
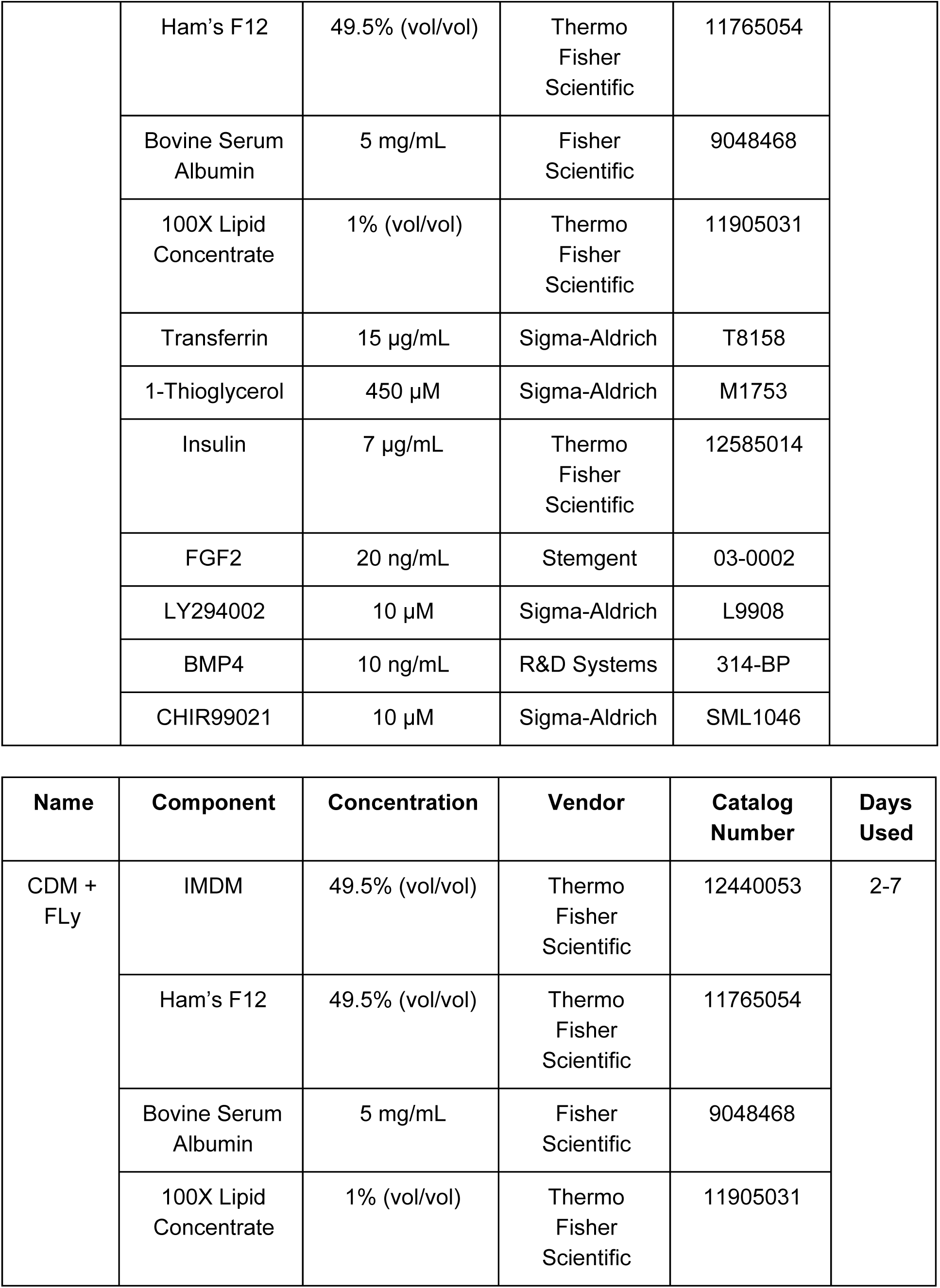
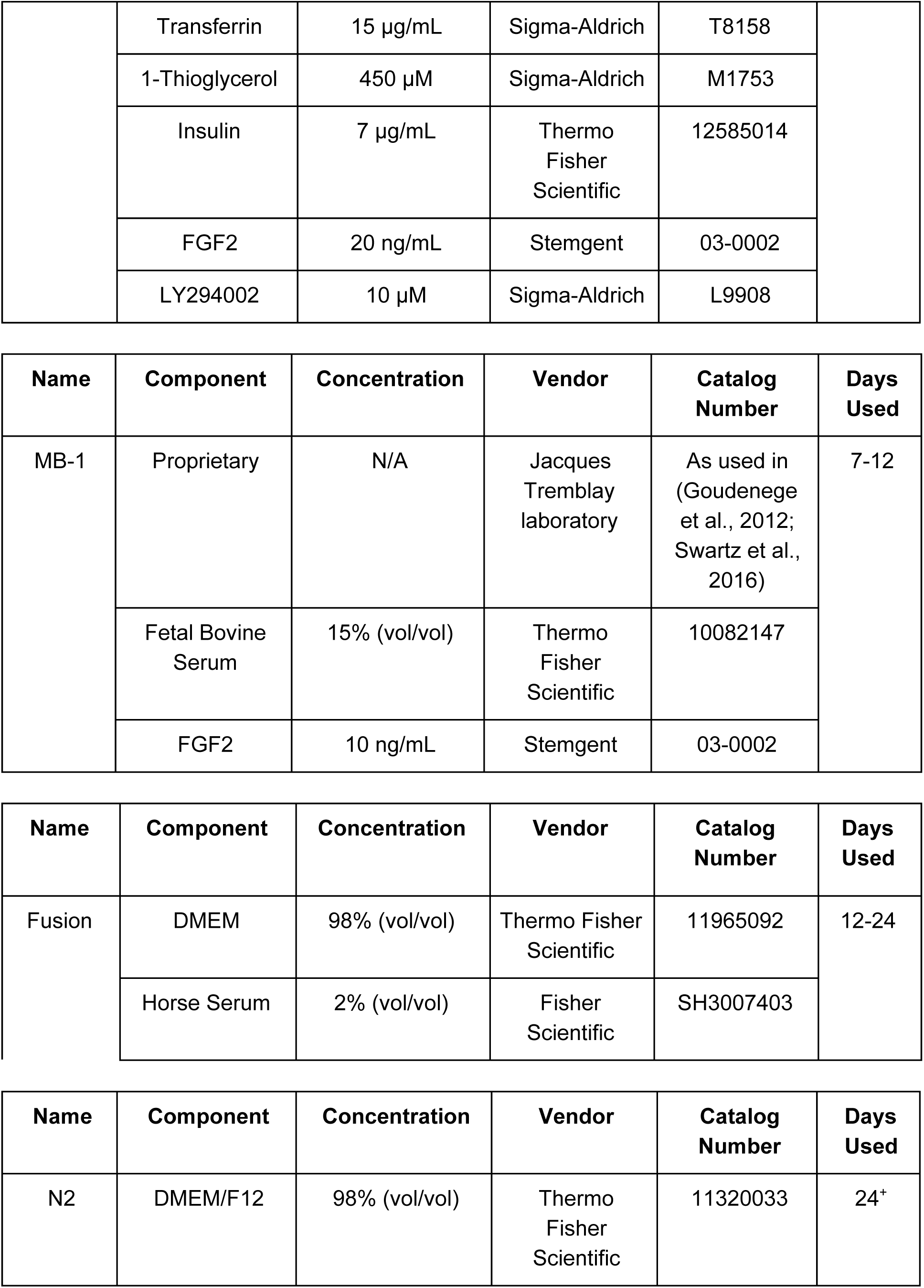
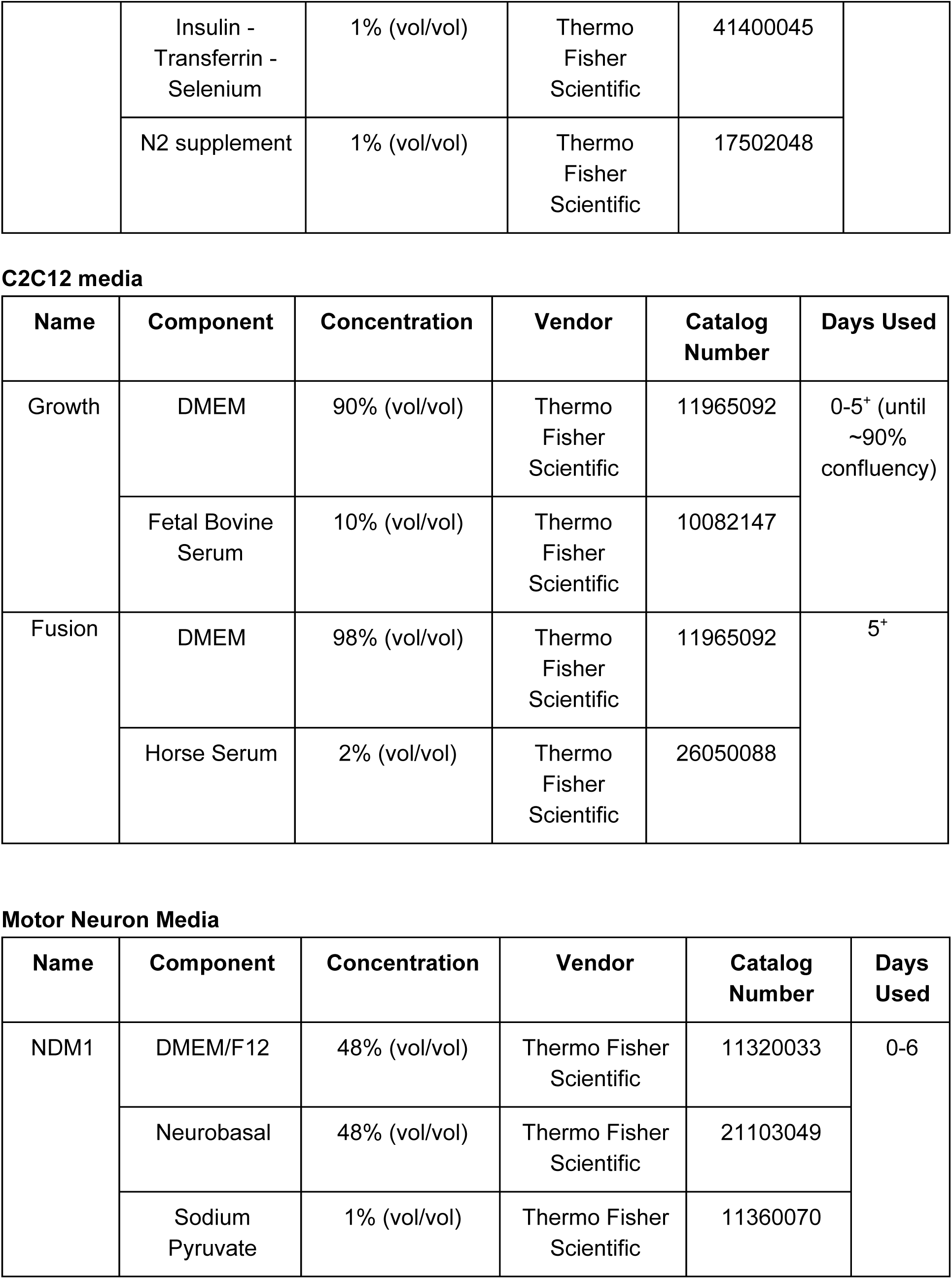
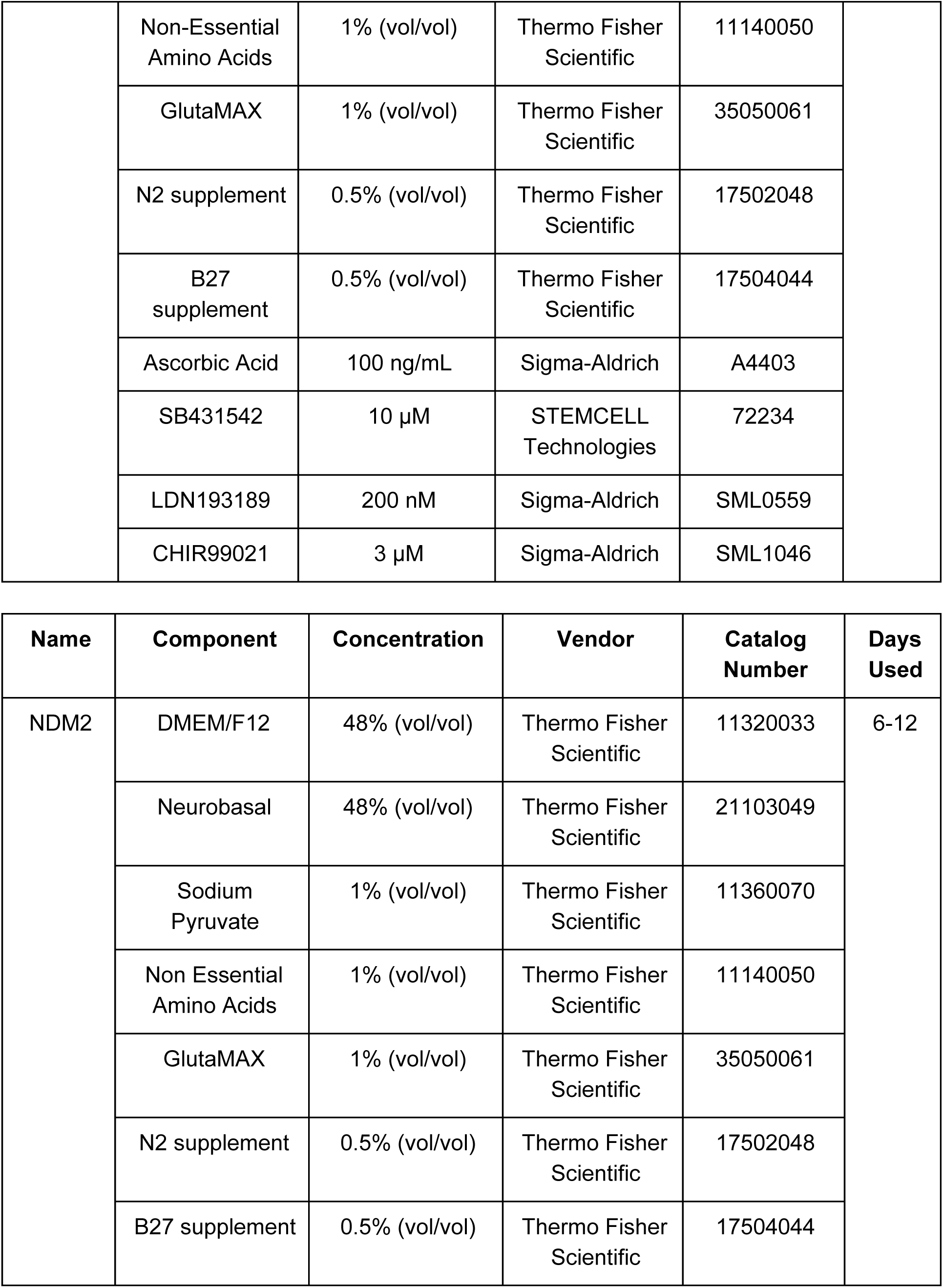
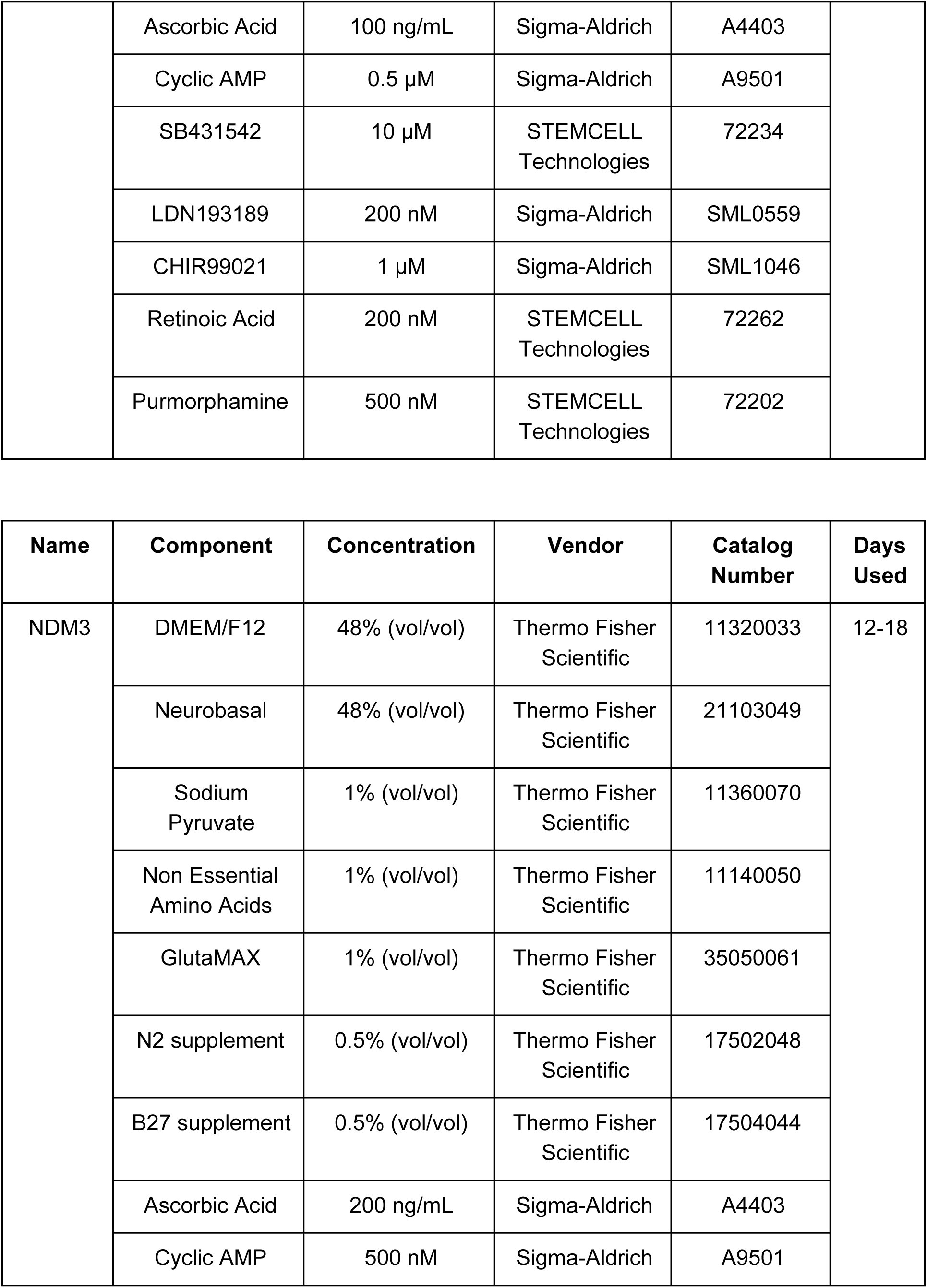
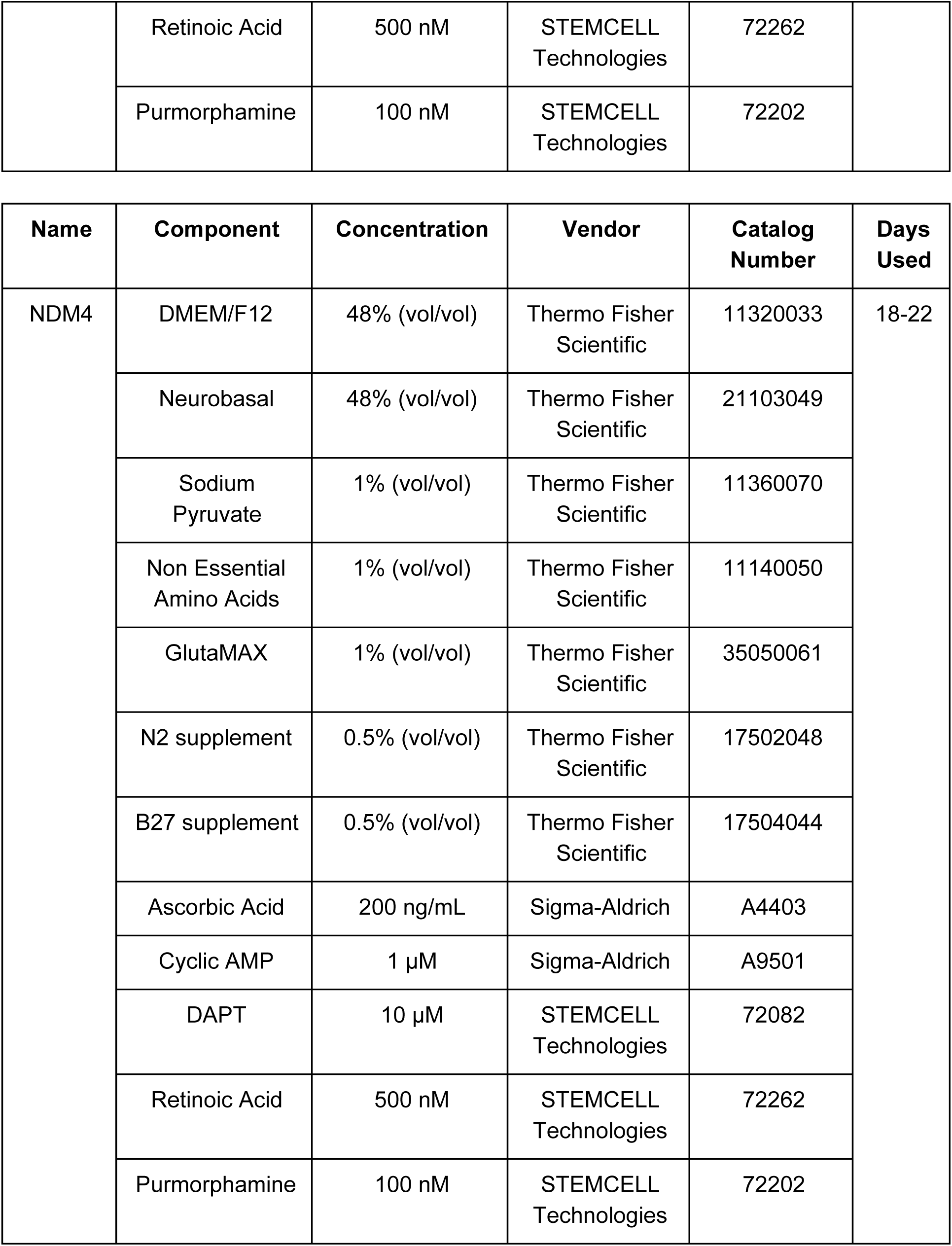
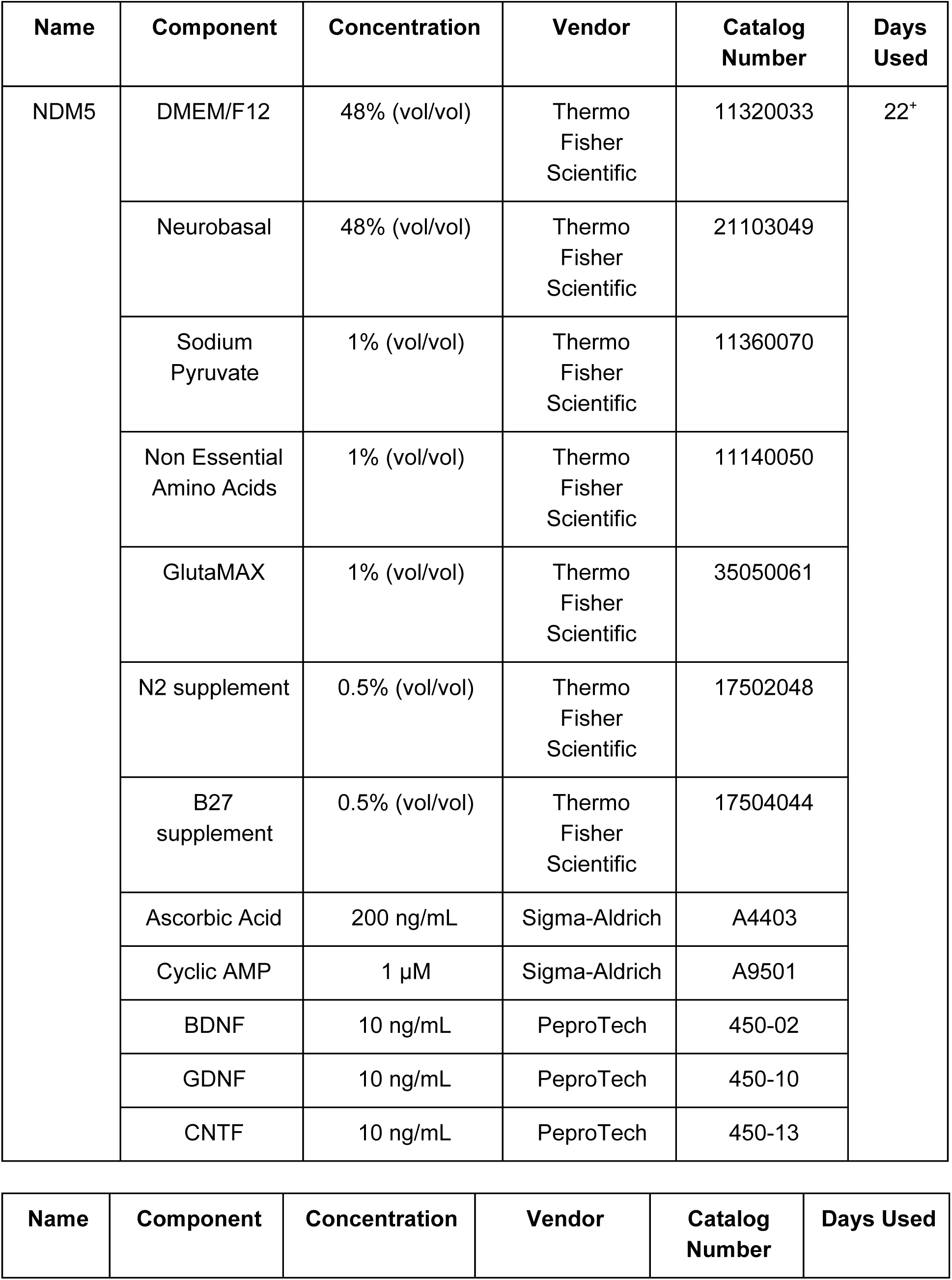
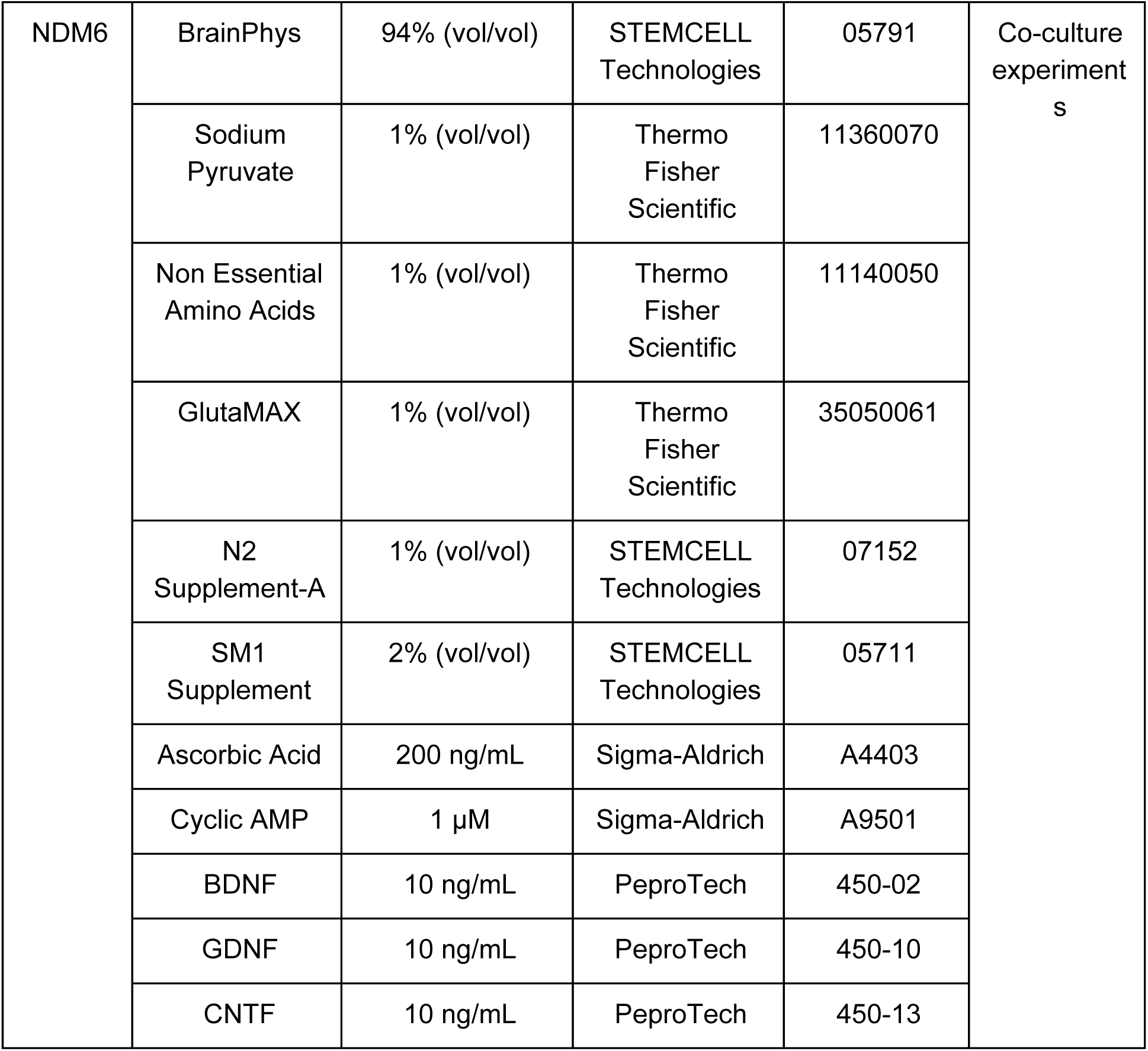
Complete list of media components used in the study. Abbreviations: DMSO, dimethylsulfoxide; FGF2, fiboblast growth factor 2; BMP4, bone morphogenic protein 4; BDNF, brain derived neurotrophic factor; GDNF, glial derived neurotrophic factor; CNTF, ciliary neurotrophic factor.

Skeletal Muscle Media
| Name | Component | Concentration | Vendor | Catalog Number | Days Used |
| --- | --- | --- | --- | --- | --- |
| TeSR-E<br>8 +<br>1.5%<br>DMSO | TeSR-E8 | 98.5% (vol/vol) | Stem Cell Technologies | 05940 | -1 to 0 |
|  | DMSO | 1.5% (vol/vol) | Sigma-Aldrich | 472301 |  |
| CDM +<br>FLyBC | IMDM | 49.5% (vol/vol) | Thermo Fisher Scientific | 12440053 | 0-2 |

|  |  |  |  |  |
| --- | --- | --- | --- | --- |
|  | Ham's F12 | 49.5% (vol/vol) | Thermo Fisher Scientific | 11765054 |
|  | Bovine Serum Albumin | 5 mg/mL | Fisher Scientific | 9048468 |
|  | 100X Lipid Concentrate | 1% (vol/vol) | Thermo Fisher Scientific | 11905031 |
|  | Transferrin | 15 µg/mL | Sigma-Aldrich | T8158 |
|  | 1-Thioglycerol | 450 µM | Sigma-Aldrich | M1753 |
|  | Insulin | 7 µg/mL | Thermo Fisher Scientific | 12585014 |
|  | FGF2 | 20 ng/mL | Stemgent | 03-0002 |
|  | LY294002 | 10 µM | Sigma-Aldrich | L9908 |
|  | BMP4 | 10 ng/mL | R&D Systems | 314-BP |
|  | CHIR99021 | 10 µM | Sigma-Aldrich | SML1046 |

| Name | Component | Concentration | Vendor | Catalog Number | Days Used |
| --- | --- | --- | --- | --- | --- |
| CDM + FLY | IMDM | 49.5% (vol/vol) | Thermo Fisher Scientific | 12440053 | 2-7 |
|  | Ham's F12 | 49.5% (vol/vol) | Thermo Fisher Scientific | 11765054 |  |
|  | Bovine Serum Albumin | 5 mg/mL | Fisher Scientific | 9048468 |  |
|  | 100X Lipid Concentrate | 1% (vol/vol) | Thermo Fisher Scientific | 11905031 |  |
|  | Transferrin | 15 µg/mL | Sigma-Aldrich | T8158 |  |
|  | 1-Thioglycerol | 450 µM | Sigma-Aldrich | M1753 |  |
|  | Insulin | 7 µg/mL | Thermo Fisher Scientific | 12585014 |  |
|  | FGF2 | 20 ng/mL | Stemgent | 03-0002 |  |
|  | LY294002 | 10 µM | Sigma-Aldrich | L9908 |  |
| MB-1 | Proprietary | N/A | Jacques Tremblay laboratory | As used in (Goudenege et al., 2012; Swartz et al., 2016) | 7-12 |
|  | Fetal Bovine Serum | 15% (vol/vol) | Thermo Fisher Scientific | 10082147 |  |
|  | FGF2 | 10 ng/mL | Stemgent | 03-0002 |  |
| Fusion | DMEM | 98% (vol/vol) | Thermo Fisher Scientific | 11965092 | 12-24 |
|  | Horse Serum | 2% (vol/vol) | Fisher Scientific | SH3007403 |  |
| N2 | DMEM/F12 | 98% (vol/vol) | Thermo Fisher Scientific | 11320033 | 24<sup>+</sup> |
|  | Insulin -<br>Transferrin -<br>Selenium | 1% (vol/vol) | Thermo<br>Fisher<br>Scientific | 41400045 |  |
|  | N2 supplement | 1% (vol/vol) | Thermo<br>Fisher<br>Scientific | 17502048 |  |

C2C12 media
| Name | Component | Concentration | Vendor | Catalog Number | Days Used |
| --- | --- | --- | --- | --- | --- |
| Growth | DMEM | 90% (vol/vol) | Thermo<br>Fisher<br>Scientific | 11965092 | 0-5<sup>+</sup> (until<br>~90%<br>confluency) |
|  | Fetal Bovine<br>Serum | 10% (vol/vol) | Thermo<br>Fisher<br>Scientific | 10082147 |  |
| Fusion | DMEM | 98% (vol/vol) | Thermo<br>Fisher<br>Scientific | 11965092 | 5<sup>+</sup> |
|  | Horse Serum | 2% (vol/vol) | Thermo<br>Fisher<br>Scientific | 26050088 |  |

| Name | Component | Concentration | Vendor | Catalog Number | Days Used |
| --- | --- | --- | --- | --- | --- |
| NDM1 | DMEM/F12 | 48% (vol/vol) | Thermo Fisher<br>Scientific | 11320033 | 0-6 |
|  | Neurobasal | 48% (vol/vol) | Thermo Fisher<br>Scientific | 21103049 |  |
|  | Sodium<br>Pyruvate | 1% (vol/vol) | Thermo Fisher<br>Scientific | 11360070 |  |
|  | Non-Essential Amino Acids | 1% (vol/vol) | Thermo Fisher Scientific | 11140050 |  |
|  | GlutaMAX | 1% (vol/vol) | Thermo Fisher Scientific | 35050061 |  |
|  | N2 supplement | 0.5% (vol/vol) | Thermo Fisher Scientific | 17502048 |  |
|  | B27 supplement | 0.5% (vol/vol) | Thermo Fisher Scientific | 17504044 |  |
|  | Ascorbic Acid | 100 ng/mL | Sigma-Aldrich | A4403 |  |
| | SB431542 | 10 μM | STEMCELL Technologies | 72234 | |
|  | LDN193189 | 200 nM | Sigma-Aldrich | SML0559 |  |
| | CHIR99021 | 3 μM | Sigma-Aldrich | SML1046 | |
| NDM2 | DMEM/F12 | 48% (vol/vol) | Thermo Fisher Scientific | 11320033 | 6-12 |
|  | Neurobasal | 48% (vol/vol) | Thermo Fisher Scientific | 21103049 |  |
|  | Sodium Pyruvate | 1% (vol/vol) | Thermo Fisher Scientific | 11360070 |  |
|  | Non Essential Amino Acids | 1% (vol/vol) | Thermo Fisher Scientific | 11140050 |  |
|  | GlutaMAX | 1% (vol/vol) | Thermo Fisher Scientific | 35050061 |  |
|  | N2 supplement | 0.5% (vol/vol) | Thermo Fisher Scientific | 17502048 |  |
|  | B27 supplement | 0.5% (vol/vol) | Thermo Fisher Scientific | 17504044 |  |
|  | Ascorbic Acid | 100 ng/mL | Sigma-Aldrich | A4403 |  |
|  | Cyclic AMP | 0.5 µM | Sigma-Aldrich | A9501 |  |
|  | SB431542 | 10 µM | STEMCELL Technologies | 72234 |  |
|  | LDN193189 | 200 nM | Sigma-Aldrich | SML0559 |  |
|  | CHIR99021 | 1 µM | Sigma-Aldrich | SML1046 |  |
|  | Retinoic Acid | 200 nM | STEMCELL Technologies | 72262 |  |
|  | Purmorphamine | 500 nM | STEMCELL Technologies | 72202 |  |
| NDM3 | DMEM/F12 | 48% (vol/vol) | Thermo Fisher Scientific | 11320033 | 12-18 |
|  | Neurobasal | 48% (vol/vol) | Thermo Fisher Scientific | 21103049 |  |
|  | Sodium Pyruvate | 1% (vol/vol) | Thermo Fisher Scientific | 11360070 |  |
|  | Non Essential Amino Acids | 1% (vol/vol) | Thermo Fisher Scientific | 11140050 |  |
|  | GlutaMAX | 1% (vol/vol) | Thermo Fisher Scientific | 35050061 |  |
|  | N2 supplement | 0.5% (vol/vol) | Thermo Fisher Scientific | 17502048 |  |
|  | B27 supplement | 0.5% (vol/vol) | Thermo Fisher Scientific | 17504044 |  |
|  | Ascorbic Acid | 200 ng/mL | Sigma-Aldrich | A4403 |  |
|  | Cyclic AMP | 500 nM | Sigma-Aldrich | A9501 |  |
|  | Retinoic Acid | 500 nM | STEMCELL Technologies | 72262 |  |
|  | Purmorphamine | 100 nM | STEMCELL Technologies | 72202 |  |
| NDM4 | DMEM/F12 | 48% (vol/vol) | Thermo Fisher Scientific | 11320033 | 18-22 |
|  | Neurobasal | 48% (vol/vol) | Thermo Fisher Scientific | 21103049 |  |
|  | Sodium Pyruvate | 1% (vol/vol) | Thermo Fisher Scientific | 11360070 |  |
|  | Non Essential Amino Acids | 1% (vol/vol) | Thermo Fisher Scientific | 11140050 |  |
|  | GlutaMAX | 1% (vol/vol) | Thermo Fisher Scientific | 35050061 |  |
|  | N2 supplement | 0.5% (vol/vol) | Thermo Fisher Scientific | 17502048 |  |
|  | B27 supplement | 0.5% (vol/vol) | Thermo Fisher Scientific | 17504044 |  |
|  | Ascorbic Acid | 200 ng/mL | Sigma-Aldrich | A4403 |  |
| | Cyclic AMP | 1 μM | Sigma-Aldrich | A9501 | |
| | DAPT | 10 μM | STEMCELL Technologies | 72082 | |
|  | Retinoic Acid | 500 nM | STEMCELL Technologies | 72262 |  |
|  | Purmorphamine | 100 nM | STEMCELL Technologies | 72202 |  |
| NDM5 | DMEM/F12 | 48% (vol/vol) | Thermo Fisher Scientific | 11320033 | 22<sup>+</sup> |
|  | Neurobasal | 48% (vol/vol) | Thermo Fisher Scientific | 21103049 |  |
|  | Sodium Pyruvate | 1% (vol/vol) | Thermo Fisher Scientific | 11360070 |  |
|  | Non Essential Amino Acids | 1% (vol/vol) | Thermo Fisher Scientific | 11140050 |  |
|  | GlutaMAX | 1% (vol/vol) | Thermo Fisher Scientific | 35050061 |  |
|  | N2 supplement | 0.5% (vol/vol) | Thermo Fisher Scientific | 17502048 |  |
|  | B27 supplement | 0.5% (vol/vol) | Thermo Fisher Scientific | 17504044 |  |
|  | Ascorbic Acid | 200 ng/mL | Sigma-Aldrich | A4403 |  |
| | Cyclic AMP | 1 μM | Sigma-Aldrich | A9501 | |
|  | BDNF | 10 ng/mL | PeproTech | 450-02 |  |
|  | GDNF | 10 ng/mL | PeproTech | 450-10 |  |
|  | CNTF | 10 ng/mL | PeproTech | 450-13 |  |
| NDM6 | BrainPhys | 94% (vol/vol) | STEMCELL Technologies | 05791 | Co-culture experiments |
|  | Sodium Pyruvate | 1% (vol/vol) | Thermo Fisher Scientific | 11360070 |  |
|  | Non Essential Amino Acids | 1% (vol/vol) | Thermo Fisher Scientific | 11140050 |  |
|  | GlutaMAX | 1% (vol/vol) | Thermo Fisher Scientific | 35050061 |  |
|  | N2 Supplement-A | 1% (vol/vol) | STEMCELL Technologies | 07152 |  |
|  | SM1 Supplement | 2% (vol/vol) | STEMCELL Technologies | 05711 |  |
|  | Ascorbic Acid | 200 ng/mL | Sigma-Aldrich | A4403 |  |
| | Cyclic AMP | 1 μM | Sigma-Aldrich | A9501 | |
|  | BDNF | 10 ng/mL | PeproTech | 450-02 |  |
|  | GDNF | 10 ng/mL | PeproTech | 450-10 |  |
|  | CNTF | 10 ng/mL | PeproTech | 450-13 |  |

### Cryopreservation of iPSC-derived Cells

All iPSCs and iPSC-derived cell vials were cryopreserved in a 1:1 mixture of the medium in which they were growing in at the time of freezing combined with a filtered 80% FBS, 20% DMSO freezing medium in a total of 1 mL per vial. Vials were immediately moved to a Mr. Frosty (ThermoFisher) container and stored at -80°C overnight before being moved to liquid nitrogen for long-term storage.

### Creation of Stable, Optogenetic iPSC lines

Isogenic *C9orf72* G_4_C_2_ carrier patient-derived lines were genome engineered using CRISPR/Cas9 as previously described (Pribadi et al., 2016). To isolate lines stably expressing optogenetic constructs, an approximately 50% confluent, single well of iPSCs (6 well format) were pre-treated with polybrene (6 µg/mL, Sigma) for 15 minutes, followed by 10 µM of ROCK inhibitor (Y-27632, Stemgent) for 60 minutes. iPSCs were dissociated to single cells following 20-25 minutes of Accutase treatment, centrifuged at 400 x *g* for 5 minutes, and re-suspended in 200µL of TeSR-E8 medium with 10 µM ROCK inhibitor in a 1.5 mL microcentrifuge tube. Two lentiviral constructs (cloned and generously donated by Dr. Ken Yamauchi, Novitch lab, UCLA) were co-transduced by adding 15 µL of each virus (>1×10^8^ titer) to the re-suspended cells. The tube was incubated at 37°C and 5% CO_2_ for 6 hours, re-suspending the cells by flicking each hour.

After 6 hours, cells were then gently re-suspended by pipetting and plated in TeSR-E8 with 10 µM ROCK inhibitor at single cell density in 48 well plates coated in vitronectin. Resulting colonies were allowed to grow for ∼7-10 days. Wells containing a single clonal population were marked and colonies were manually passaged using mechanical dissociation with a P20 pipette tip and transferred to a new plate. Residual cells from the colony were lysed and screened by PCR for *Cre* and *ChannelRhodopsin 2* (*ChR2*) genomic DNA integration. 22 total double-positive clones were differentiated in parallel to motor neurons. 5/11 clones from each line were selected based on YFP fluorescent strength on day 30 and a subset of these were screened by qPCR for expression of *Cre* and *ChR2*. 1 clone from each line with the highest transgene expression was chosen for further study and the G_4_C_2_ repeat status was confirmed by repeat-primed PCR.

### Cell Lysis and PCR Screening

Cells were lysed *in situ* within 48 well plates. 100 uL of Lysis Buffer (10% 10X AccuPrime PCR Buffer I + 200 µg/mL Proteinase K in H_2_0) was added to each well. Cells were then incubated at 60°C for 10 minutes to assist with the lysis. Cells were then further broken down manually via gentle pipetting. Lysates were thereafter vortexed, transferred to PCR plates, and incubated overnight at 60°C. The next day, plates were centrifuged briefly to collect lysate at the bottom of each well and then heated at 95°C for 20 minutes to deactivate the Proteinase K.

### Neuromuscular Junction Area and Myotube Width Calculations

Vials from previously differentiated iPSC-sKM (line isoA80) frozen on day 79 of differentiation were thawed onto 12 mm coverslips coated in Matrigel in 24 well format as previously described. Skeletal myotubes were allowed to grow in N2 medium for 3 days. On day 3, coverslips were switched to 3 conditions: (1) in N2 medium, (2) in a 1:1 mixture of NDM5 and NDM6 mediums containing recombinant human Agrin (50 ng/mL), or (3) in a 1:1 mixture of NDM5 and NDM6 mediums containing recombinant human Agrin (50 ng/mL) and a single MN sphere. For condition (3), a single, day 40 iPSC-MN sphere (Opto-A1, Opto-isoA80-8) was plated onto each coverslip on Day 3 and cultured using methods described previously. A single sphere was utilized to limit variability in culture conditions.

Starting on Day 3, each coverslip was imaged every 48 hours in several random fields of view until Day 9 (Day 6 of co-culture) to determine change in myotube width over time in each condition. Images were then scored manually using Q-Capture Pro 7 software. 5 multinucleated myotubes per image were scored by measuring the thickest width of the myotube. At least 5 individual coverslips contributed to each condition.

For condition (3), cells were fixed as described below (Immunocytochemistry) on Day 10 (Day 7 of co-culture) and stained with Alexa-fluor 594-conjugated α-Bungarotoxin (α-BTX, 1:500) for 45 minutes at room temperature. Coverslips were then washed 3x in PBS, transferred to 35 mm round-bottom glass dishes (MatTek), and imaged on a spinning disc confocal microscope as described below (Optogenetically-evoked Calcium Recordings). Regions around the periphery of the MN sphere showing co-localized α-BTX and YFP were imaged as 10-20 µm Z-stack sections at 20X magnification. At least 5 coverslips from each line (Opto-A1, Opto-isoA80-8) were imaged, totaling >25 images for each condition. Images were randomized and scored by a blinded observer using the native Draw Spline Contour tool in ZENBlue software. Only regions displaying α-BTX and YFP co-localization were scored and areas below 5 µm^2^ were excluded from analysis as being potentially indistinguishable from background.

### Optogenetically-evoked Calcium Recordings

All co-culture optogenetic experiments were performed on Day 7 using MN spheres between days 40-60 *in vitro* on 35 mm glass bottom dishes (MatTek, catalog # P35G-1.5-20-C). The age of myotubes included in experiments varied between days 60-100 *in vitro* at the time of data collection. MN spheres were derived from Opto-A1 and Opto-isoA80-8 whereas all myotubes were derived from isoA80.

Experiments were performed on a Zeiss Axio Observer Z1 spinning disc confocal microscope outfit with an EMCCD camera and incubation system at 37°C and 5% CO_2_. The microscope was outfit with a blue LED (ThorLabs M470L2, 470nm) and custom filter cube (Chroma) to reflect wavelengths below 488 nm and pass wavelengths above 561 nm. The LED light stimulation settings were controlled by a Master-8 pulse stimulator.

Two hours prior to recording, conditioned medium from co-cultures to be recorded from was collected. Co-cultures were incubated with the cell permeable calcium dye X-Rhod-1 AM (2 µM, Life Technologies) for 25 minutes at 37°C in fresh NDM6 containing recombinant human Agrin (50 ng/mL), washed 3x for 5 minutes in NDM6 and then cultured for at least 30 minutes in the conditioned medium to allow for de-esterification of the dye prior to recording.

Data was collected using ZENBlue software (Zeiss) in 20 second videos where the stimulation paradigm consisted of five 250 ms LED pulses at 1 Hz. Regions of interest (ROIs) were manually selected on individual myotubes and normalized to a background ROI containing the same pixel volume (4.5 pixel diameter circle). ΔF/F ratiometric measurements were performed using ZENBlue software, where F_t0_ was calculated from the average intensity across the total number of frames from the whole experiment. Intensity and ratiometric plots were generated in RStudio (version 1.1.383) from the exported raw data.

### Optogenetics in iPSC-sKM

Lentiviral particles encoding a ChR2 (H134R)-eYFP fusion protein (vector ID: VB170830-1111twn) were packaged by VectorBuilder. iPSC-sKM (> day 50) were treated with 5 µg/mL of polybrene for 20 minutes, washed twice in N2 medium, and transduced with 10 µL of virus (titer 2.14×10^8^) for 24 hours. iPSC-sKM was then grown in N2 medium for up to 4 weeks prior to recording. Live calcium imaging was performed as previously described (Optogenetically-evoked Calcium Recordings), using the same cell permeable calcium dye X-Rhod-1 AM, data analysis methods, with multiple different light stimulation protocol attempts.

### Image and Video Acquisition

Images (1F-H, 1K, 2D) were taken on a Zeiss LSM 310 confocal microscope and z-stacks were merged using ImageJ software. Images (3E-F, 3J) were taken on a Zeiss Axio Observer Z1 spinning disc confocal microscope.

All other images and videos were taken on an Olympus iX53 microscope outfit with a Retiga EXi Fast 1394 camera using Q-Capture Pro 7 software. Videos were taken with maximum frame speed enabled and converted in ImageJ at 7 frames per second. Videos were stitched or edited using iMovie software.

### Immunocytochemistry

Cells were fixed in cold 4% paraformaldehyde (PFA) for 15 minutes, rinsed 3x for 5 minutes in phosphate-buffered saline (PBS), and stored at 4°C until staining. Cells were permeabilized in buffer (0.1% Triton-X, 2.5% donkey serum in PBST (0.1% Tween-20 in PBS)) for 1 hour at room temperature (RT). Primary antibody was added in permeabilization buffer overnight at 4°C on a rocker on a low setting to prevent detachment of cells. The next day, cells were rinsed 2x with PBS for 10 minutes and Alexa Fluor-conjugated secondary antibodies were added in 0.1% Triton-X and 5% donkey serum in PBS for 1 hour at RT in the dark. Cells were rinsed 2x with PBS for 10 minutes and DAPI or TOPRO-3, if necessary, was added at 1:1000 in PBS for 5 minutes at RT, washed 2x in PBS for 10 minutes, and stored in the dark until imaging. Coverslips were mounted in ProLong gold and allowed to set overnight at 4°C prior to imaging. A complete list of antibodies used can be found in **Table S3**.

**Table S3.** List of antibodies used in the study. DSHB, Developmental Studies Hybridoma Bank.

| **Gene** | **Cat. Number** | **Vendor** | **Host Species** |
| --- | --- | --- | --- |
| Anti-mouse AP2α, monoclonal | DSHB | 3B5 | 1:20 |
| Anti-goat ChAT, polyclonal | Millipore | AB144P | 1:300 |
| Anti-rabbit Connexin43, polyclonal | Abcam | ab11370 | 1:500 |
| Anti-mouse HB9, monoclonal | DSHB | 81.5C10 | 1:20 |
| Anti-mouse HNK1, monoclonal | DSHB | 3H5 | 1:15 |
| Anti-mouse ISL1, monoclonal | GeneTex | GTX83120 | 1:200 |
| Anti-rabbit Myelin Protein Zero (MPZ), polyclonal | Abcam | ab31851 | 1:150 |
| Anti-mouse Myosin Heavy Chain (MHC), Monoclonal | DSHB | MF-20 | 1:40 |
| Anti-rabbit OLIG2, polyclonal | Millipore | AB9610 | 1:200 |
| Anti-mouse Synaptic Vesicle 2a (SV2A), monoclonal | DSHB | SV2 | 1:30 |
| Anti-mouse TITIN, monoclonal | DSHB | 9-D10 | 1:40 |
| Anti-rabbit Beta-3-Tubulin (TUJ1), polyclonal | Covance | PRB-435P | 1:1000 |
| Anti-goat ChAT, polyclonal | DSHB | 3B5 | 1:20 |

### Patch-clamp electrophysiology

Standard whole-cell patch clamp methods were applied to record currents from muscle cells with an Axopatch 200B amplifier and a Digidata 1321A at room temperature in a bath solution containing (in mM) 140 NaCl, 2.5 KCl, 2 CaCl2, 2 MgCl2, 10 glucose, and 10 HEPES (pH 7.3). The intrapipette solution contained 150 KCl, 1 NaCl, 1 MgCl2, 5 EGTA, and 10 HEPES (pH 7.3). Glass pipettes were made from patch glass with 1.5 mm O.D. and 1.17 mm I.D. Multi-step pulling was done with a Sutter P-97 micropipette puller and the pipette resistance ranged from 2-5 MW when filled with intrapipette solution. Cells were held at -70 mV with 70-80% compensation of series resistance. Data were acquired and analyzed using pClamp9 software. Data were sampled at 10 KHz and filtered at 1-2 kHz. Graphs were plotted using Origin software.

### Gap Junction Blockade

iPSC-sKM were grown to confluence and observed periodically for spontaneous contractions. Baseline spontaneous contractions were recorded for 20 seconds and 1-heptanol was added to the medium to a final concentration of 5 mM. 20 second recordings from the same well position were taken 5 and 10 minutes after addition of 1-heptanol. 1-heptanol was washed out twice in N2 medium and cells were allowed to equilibrate at 37°C for 15 minutes before recording again. iPSC-sKM were 48 days *in vitro* at the time of recording.

### RT-PCR, qRT-PCR, and RP-PCR

For RT-PCR, RNA was isolated from frozen cell pellets using the RNeasy Plus Mini Kit according to the manufacturer’s protocol. Complementary DNA (cDNA) synthesis was made from 500 ng total RNA using the SuperScript III First-Strand Synthesis kit. PCR was run using 10 ng of cDNA per reaction. Products were run on a gel and imaged. Total RNA from human skeletal muscle was obtained from a 20-year-old male (Clontech).

For qPCR, cDNA was obtained using the same methods. Relative gene expression levels were determined in triplicate reactions of 10 ng cDNA per reaction using the SensiFAST SyBR kit on a LightCycler 480. Quantification was performed using the –ΔΔCt method normalized to the geometric mean of *GAPDH* and *ACTB* as housekeeping genes. A complete list of primers used in the study can be found in **Tables S4** and **S5**.

For repeat-primed PCR (RP-PCR), *C9orf72* repeat expansion mutations were determined as described by DeJesus-Hernandez et al (DeJesus-Hernandez et al., 2011). PCR products were run on an ABI3730 DNA Analyzer and analyzed using Peak Scanner software. The characteristic sawtooth pattern is indicative of the presence of a repeat expansion.

**Table S4.**
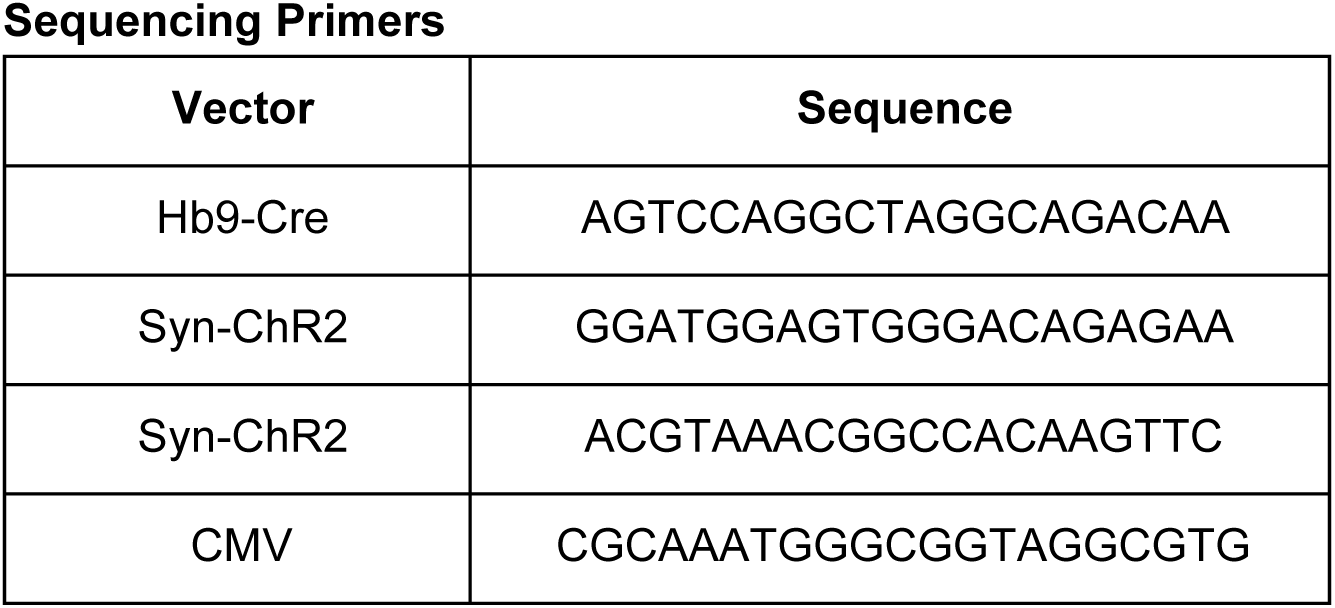
Sequencing primers used for confirmation of viral vectors.

**Table S5.**
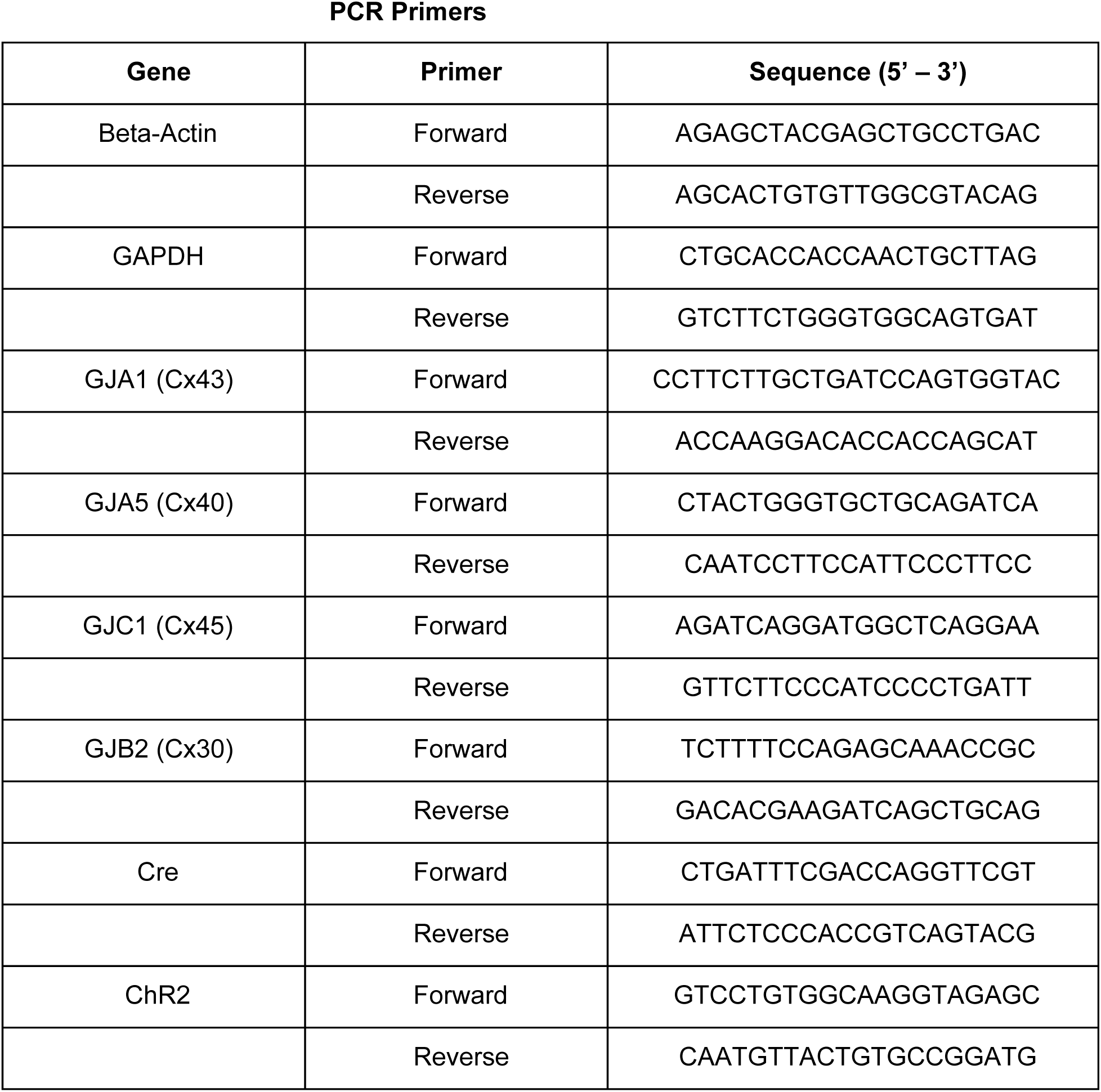
Primer sets used for PCR experiments in this study.

### Electron Microscopy

iPSC-sKM were grown on Matrigel-coated Thermanox coverslips. The cultures were then fixed for 45 minutes in 4% paraformaldehyde, 1% glutaraldhedye in 0.1M phosphate buffer and washed several times in buffer. They were then post-fixed for 1 hour in 1% OsO_4_ in ddH_2_O at RT, washed 3x in ddH_2_O, and dehydrated through a graded series of ethanols. They were briefly treated with propylene oxide before infiltration with a 1:1 mixture of Eponate 12 and propylene oxide for 1 hour followed by a 2:1 mixture overnight. The caps were slit to allow the propylene oxide to slowly evaporate. The following day, the cells were placed in fresh Eponate, placed under vacuum for several hours and polymerized at 60C for 48 hours. Silver sections were cut on a RMC MTX ultramicrotome and placed on formvar-coated slot grids. The grids were stained with saturated uranyl acetate and Reynold’s lead citrate before examination on a JEOL 100CX electron microscope at 60kV. Images were collected on film, and the negatives scanned at 1200 dpi.

### Multi-Electrode Arrays

For pharmacological experiments, a previously frozen vial of iPSC-sKM (day 79, isoA80) was thawed and plated onto a 48 well multi-electrode array plate (tMEA-48OPT, Axion Biosystems). iPSC-sKM was grown for 3 days in N2 medium and on day 3, live iPSC-MN spheres (day 40, Opto-isoA80-8) were plated on top, creating co-cultures. Co-cultures were grown in NDM5 and ¾ medium was replaced every 48 hours with NDM6 containing recombinant human Agrin (50 ng/µL). At the time of recording, the co-culture was 7 days old, with the last medium change occurring 48 hours prior.

Electrophysiological activity was recorded using the Maestro Pro (Axion Biosystems) at 37°C. Data was acquired as raw voltage using a bandpass filter (200 – 4000 Hz) and a gain of 1000x at a sampling rate of 12.5 kHz. Neural spikes were identified as peaks in voltage surpassing 6 timex the standard deviation on each electrode. Spike data was processed further and statistics generated using AxIS Navigator and Axion Neural Metric Tool (Axion Biosystems). Optogenetic stimulation was performed with the Lumos (Axion Biosystems) using 475 nm light at 50% power with 500 ms pulses at 0.25 Hz. Wells were prioritized based on qualitative assessment of their spontaneous activity and optogenetic response. Prior to drug treatment, baseline activity was recorded for 5 minutes from spontaneous and optogenetically-paced samples. For wells to be treated with vecuronium bromide or decamethonium bromide, vehicle (0.2% DMSO) was added to each chosen well and the plate was incubated for 20 minutes at 37°C and 5%CO_2_. For 1-heptanol, NDM6 was used as the solvent. The plate was then returned to the Maestro Pro device and allowed to equilibrate for 2 minutes before recording optogenetically-evoked activity for 5 minutes. This protocol was then repeated for each drug treatment, using the same well that previously received vehicle. Drug concentrations were informed by previous use in the literature, followed by 10 fold decreases.

Gross functional activity was quantified as the number of spikes during a particular recording. Synchronous well-wide activity induced via light stimulation was quantified as the network burst frequency and the percentage of spikes that belong to a network burst. Response to drug treatment in optogenetically-paced cultures was quantified by the average number of spikes in a 500 ms window post-stimulation only on electrodes yielding an evoked response.

### Statistics

Statistics were performed in RStudio (version 1.1.383). Collected sample datasets were tested for normality using the Shapiro-Wilk test. For muscle width calculations, a 1-way ANOVA test was performed, followed by pairwise-t tests with Bonferroni correction for multiple comparisons and confirmed with Tukey’s HSD post-hoc test. For NMJ area calculations, the non-parametric data were resampled 15,000 times using bootstrapping methodology (Curran-Everett, 2009) where a p-value was obtained via a null hypothesis significance test conducted on differences in the medians between test groups. Effect magnitude was determined by re-sampling median differences 15,000 times and reporting the resultant 95% confidence intervals on the sorted median differences. For MEA data, Wilcoxon rank sum tests were performed. Statistical tests were not performed on datasets with no variance.

